# Severe and multifaceted systemic immunosuppression caused by experimental cancers of the central nervous system requires release of non-steroid soluble mediators

**DOI:** 10.1101/2020.03.24.006825

**Authors:** K Ayasoufi, CK Pfaller, L Evgin, RH Khadka, ZP Tritz, EN Goddery, CE Fain, LT Yokanovich, BT Himes, F Jin, J Zheng, MR Schuelke, MJ Hansen, W Tung, LR Pease, RG Vile, AJ Johnson

## Abstract

Immunosuppression of unknown etiology is a hallmark feature of glioblastoma (GBM) and is characterized by decreased CD4 T cell counts and down regulation of MHC class II expression on peripheral blood monocytes in patients. This immunosuppression is a critical barrier to the successful development of immunotherapies for GBM. We recapitulated the immunosuppression observed in GBM patients in the C57BL/6 mouse and investigated the etiology of low CD4 T cell counts. We determined that thymic involution was a hallmark feature of immunosuppression in three distinct models of CNS cancer, including mice harboring GL261 glioma, B16 melanoma, and in a spontaneous model of Diffuse Intrinsic Pontine Glioma (DIPG). In addition to thymic involution, we determined that tumor growth in the brain induced significant splenic involution, reductions in peripheral T cells, reduced MHC class II expression on hematopoietic cells, and a modest increase in bone marrow resident CD4 T cells with a naïve phenotype. Using parabiosis we report that thymic involution, declines in peripheral T cell counts, and reduced MHC class II expression levels were mediated through circulating blood-derived factors. Conversely, T cell sequestration in the bone marrow was not governed through circulating factors. Serum isolated from glioma-bearing mice potently inhibited proliferation and functions of T cells both *in vitro* and *in vivo*. Interestingly, the factor responsible for immunosuppression in serum is nonsteroidal and of high molecular weight. Through further analysis of neurological disease models, we determined that the aforementioned immunosuppression was not unique to cancer itself, but rather occurs in response to CNS injury. Noncancerous acute neurological insults also induced significant thymic involution and rendered serum immunosuppressive. Both thymic involution and serum-derived immunosuppression were reversible upon clearance of brain insults. These findings demonstrate that CNS cancers cause multifaceted immunosuppression and pinpoint circulating factors as a target of intervention to restore immunity.

**Short Summary:** CNS cancers and other brain-injuries suppress immunity through release of non-steroid soluble factors that disrupt immune homeostasis and dampen responses of the peripheral immune system.

Graphical Abstract

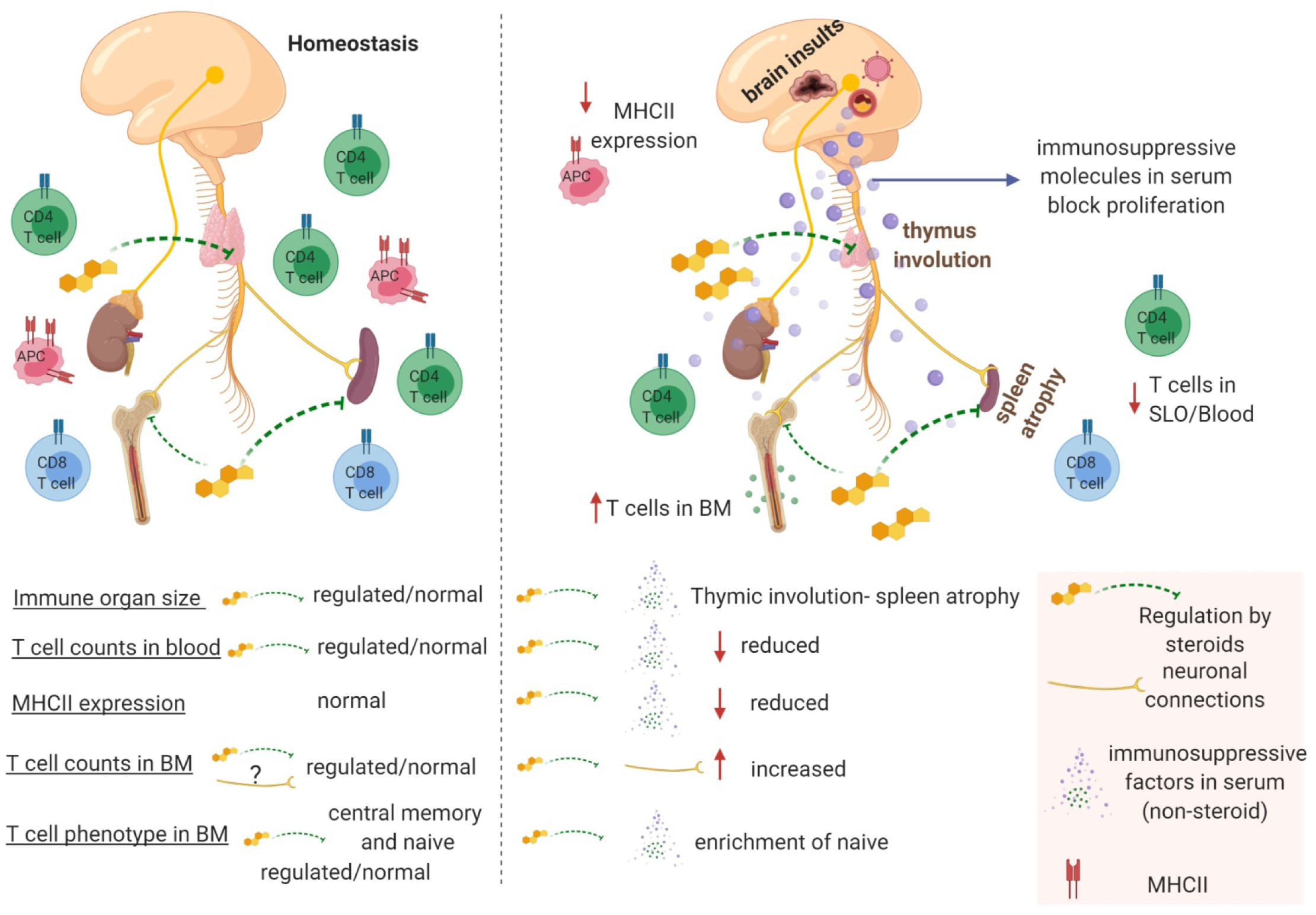

## Introduction

Systemic immunosuppression is a hallmark feature of neurological insults as diverse as stroke, traumatic brain injury, and cancers of the central nervous system (CNS) (*1–8*).This immunosuppression is a critical barrier to both patient survival and efficacy of immune modulating therapies. Glioblastoma (GBM) is an incurable malignant brain tumor with very poor prognosis that affects over 12000 patients annually in the United States alone (*9*). Hence, immunosuppression in GBM is a major barrier to the development of successful immunotherapy and viral-therapy (*2*). GBM patients can have CD4 T cell counts comparable to those with acquired immunodeficiency syndrome (AIDS), yet current efforts are focused on developing and improving immunomodulatory therapies which will be ineffective in immunosuppressed individuals (*2, 6*). While the presence of immunosuppression in neurological diseases has become an accepted feature of these conditions, the exact immunological nature and the underlying mechanisms of the immunosuppression remain largely unknown.

Peripheral immunosuppression induced by neurological insults affects both primary and secondary immune organs (*1, 10, 11*). All organs, including the thymus and bone marrow are innervated (*12–14*). While the function of organ innervation is well established for endocrine, circulatory and pulmonary systems, innervation of primary and secondary immune organs is less well-defined. A major focus of neuroimmunology research has been the activation, infiltration and induction of neuropathology by immune cells (*15–25*). However, recently there is increased focus on the bi-directionality of neuro-immune communication (*11, 26–33*). These studies indicate that the nervous system affects functions of the peripheral immune system, including bacterial clearance, aging, and allergy and itch responses through neuronal connections and/or release of soluble factors at the site of injury (*27–30, 33*). One notable example is blockade of the calcitonin gene-related peptide (CGRP) production from transient receptor potential vanilloid 1^+^ (TRPV1^+^) nociceptors which can dramatically improve bacterial clearance and neutrophil recruitment and function (*28*). Similarly, the aging process was recently linked to innervation of a primary immune organ, as degeneration of adrenergic nerves in the bone marrow was associated with premature aging in mice (*33*). These studies have immensely enhanced our understanding of communication between the peripheral nervous system and the immune system under several conditions. However, it remains unclear how neurological diseases, especially GBM, lead to immune suppression not only within the brain/tumor microenvironment, but also outside of the CNS. Studies linking injury in the CNS to a direct and measurable change in the immune system are needed because many neurological diseases cause severe immunosuppression of unknown etiology (*1, 11, 31, 32*).

The thymus is the critical primary immune organ responsible for generating and educating T cells. Additionally, the thymus is crucial in maintaining peripheral T cell counts in both children and adults (*34, 35*). As T cell development and education exclusively require a functional thymus, changes in thymic homeostasis have long lasting immunological consequences (*34, 36*). Once believed to be less important in maintaining a healthy immune system in adults, numerous studies have highlighted the crucial role of the adult thymus in generating peripheral T cells under conditions of lymphopenia (*34, 36–41*). Patients devoid of a functional thymus cannot reconstitute their T cell repertoire following hematopoietic transplantation, HIV infection, chemotherapy, and general immunosuppression (*34-36, 38-41*). Importantly, the adult thymus is still generating new T cells into the seventh decade of life, as measured by T cell Receptor Excision Circles (TRECs) quantifying recent thymic emigrants in the blood of aged adults (*36, 39*). For these reasons, we aimed to study the role of brain-thymus communication during immunosuppression induced by neurological insults such as cancers of the CNS. We hypothesized that peripheral immunosuppression following neurological insults are due to a common but complex pathway of communication between the CNS and the primary and secondary immune organs.

In this study, we investigated the extent to which the thymus has the capacity to respond to insults, including experimental GBM, which are contained within the CNS. In addition, we go beyond the thymus and evaluate the changes within spleen, peripheral blood, serum, and bone marrow of mice harboring experimental GBM. We report that the experimental-GBM-induced neuro-immune communication through the brain-thymus axis is one significant part of a multifaceted systemic immunosuppression. Our evaluation of additional neurologic disease models demonstrate that the mechanisms leading to this immunosuppression involve release of non-steroid soluble factors following brain injury. These results help us better understand changes in the immune system following CNS insults, and provide new insights into mechanisms of immunosuppression induced by CNS cancer.

## Methods

### Mice

Both male and female (WT) C57BL/6, (GFP) C57BL/6-Tg (UBC-GFP) 30Scha/J, and MHCII^-/-^ (B6-12952-^H2d1Ab1-EA^/J) mice were purchased from Jackson Laboratory (Bar Harbor, Maine). Catalog numbers are 000664, 004353, and 003584 respectively. GFP mice express GFP under the Ubiquitin C promoter which is expressed in all tissues. GFP mice and MHCII^-/-^ mice were bred in our facility. Adrenalectomized mice were purchased from Jackson Laboratory (Bar Harbor, Maine). These mice were maintained on 1% saline in drinking water for the duration of the experiments. Mice were bred and maintained in the animal facility at Mayo Clinic under Institutional Animal Care and Use Committee (IACUC) guidelines. Handling of all animals and performing of all procedures were approved by the Mayo Clinic IACUC.

### Intracranial injection

Mice were anesthetized by intraperitoneal injection of 30 mg/kg Ketamine and 3 mg/kg Xylazine. Under anesthesia, the scalp was prepared in a sterile fashion with Betadine, and a 0.5 cm longitudinal incision was made using a sterile scalpel. The skin was then retracted at the incision site and a right frontal burr hole was drilled into about 1 mm lateral and 2 mm anterior to bregma. Using a stereotactic frame, the needle of a Hamilton syringe (Hamilton Company, Reno, Nevada) was positioned in the burr hole and lowered 3.3 mm into the cortex and retracted 0.3 mm. At this depth, injections were performed in a total volume of 1.5-2µL. We injected half of the volume followed by a 2 minute pause. The second half of the volume was then injected followed by a 3 minute pause after which the needle was slowly retracted out of the brain. The wound was sutured with 4-0 vicryl (Ethicon Inc, Somerville, NJ) suture. This method was used for injection of PBS, LPS, GL261 cells, and B16 F1 melanoma cells.

### TMEV infection

For intracranial TMEV infections, mice were anesthetized with 2% isoflurane and given an intracerebral injection with 2×10^6^ PFU of Daniel’s strain of TMEV in a 10 µl volume as previously published (*17*). For intraperitoneal (IP) infection, mice were given an IP injection with 2X10^7^ PFU of Daniel’s strain of TMEV in a 100 µl volume. TMEV was made by Dr. Kevin Pavelko at Mayo Clinic (Rochester, MN) (*42, 43*).

### Vesicular stomatitis virus encoding Ovalbumin (VSV-OVA) infection and tetramer staining

This virus (VSV-OVA) was kindly provided by Dr. Richard G Vile. It was prepared in Dr. Vile’s lab as previously reported (*44*). Mice were infected with 1 x10^7^ PFU intravenously. Antigen specific responses were quantified using K^b^:OVA (SIINFEKL) tetramers. Tetramers were provided by the NIH tetramer facility (Atlanta, GA).

### Tumor cell culture (GL261-Luc, B16.F1 melanoma)

GL261-Luciferace (GL261-Luc) expressing cells were provided by the laboratory of Dr. John Ohlfest (Masonic Cancer Center University of Minnesota, Minneapolis, MN). These cells were modified to express a luciferase construct. GL261-Luc cells were grown in Dulbecco’s Modified Eagle Media (DMEM, Gibco, Gaithersburg, MD) with L-glutamine media supplemented with 10% fetal bovine serum (FBS) and 1% Penicillin/Streptomycin (Sigma, St. Louis, MO). To detach confluent cells, cells were treated with trypsin L express, (Cat # 12605-010 Gibco, Gaithersburg, MD), washed and counted. 60000 cells were resuspended in 1.5-2 µl of sterile PBS and injected into mice according to the procedure for intracranial injection.

B16.F1 melanoma cells were cultured in DMEM + 10% FBS in the laboratory of Dr. Richard Vile (Mayo Clinic, Rochester, MN). 10000 melanoma cells were implanted in 2 µL volume into the frontal lobe according to the intracranial injection procedure described above.

### Spontaneous Murine Glioma Model

RCAS-based spontaneous murine glioma system was a generous gift of Dr. Oren Becher (Northwestern University) and was used as previously described (*45*). Briefly, DF1 chicken fibroblasts cultured in DMEM with 10% FBS at 39°C and 5% CO_2_ were transfected with RCAS vectors (PDGFb, H3.3K27M, and Cre recombinase). After 2-3 weeks of passaging, 1×10^5^ DF1 cells with a 1:1:1 vector ratio or untransfected DF1 controls were implanted into the brainstems of P3-P5 Nestin Tv-a, p53^fl/fl^ pups. After 28 days, all mice underwent T2-weighted MRI imaging to assess tumor volumes. Mice with detectable tumors and non-tumor bearing control mice were euthanized for assessment of the thymus.

### Bioluminescence imaging

Tumor burden in GL261-Luciferase bearing animals was assessed using bioluminescence imaging as previously described(*19*). Animals were intraperitoneally injected with 150 mg/kg D-luciferin sodium salt in PBS (Gold Biotechnology, Olivette, MO). Animals were anesthetized with 2.5% isoflurane before imaging. 0.5-1% isoflurane was used to maintain anesthesia during imaging. Animals were scanned using Mayo Clinic’s IVIS Spectrum system (Xenogen Corp., Amameda, CA, USA) running Living Image software. Bioluminescence intensity (photons/s) was calculated by placing a circular region of interest (ROI) over the head. Several images were taken at different exposure times and averaged to generate final intensity. Mice with bioluminescence intensities above 10^5^ (photons/second, p/s) were considered to be tumor-bearing whereas those below 10^5^ (p/s) were considered to be tumor negative.

### Seizure induction

A saline Kainic acid (KA) solution was made at 2mg/ml. Mice were weighed and received an IP injection of appropriate amount of KA solution. In male mice, seizure was induced by injecting 17.5 mg/kg of KA (Sigma, St. Louis, MO) per mouse. In female mice, seizure was induced using 15.5 mg/kg of KA. To stop seizure activity at 90 minutes post KA injection, all mice received Valproic acid (Sigma, St. Louis, MO) at 300 mg/kg concentration from a solution of 45 mg/ml made in saline. We used a modified Racine score to measure seizure activity as previously described (*46, 47*). This scoring system is as follows: **Stage 0**: Normal Activity, **Stage 1**: Immobility, **Stage 2**: Rigid posture, outstretched limbs and raised tail, **Stage 3**: Seizure onset with rearing (standing on hind limbs) and forelimb clonus (shaking), **Stage 4**: Rearing and falling with continuous forepaw shaking, **Stage 5**: Continuous stage 4 often resulting in falling on side and forelimb and hind limb shaking, **Stage 6**: Discrete and intense seizure activity including jumping, and **Stage 7**: Death. Mice that had at least progressed through stage 1 were included in the study. Animals with scores of 0 were excluded from the study. 50% of males experienced a stage 4/5 seizure and died within the first hour post KA injection, thus only the remaining mice were used for analysis. Female mice that did not have any seizure activity by 50 minutes were injected with an extra 3 mg/kg of KA and their seizure activity was monitored.

### Organ processing

Spleens and thymi were dissected and weighed before dissociation using the rubber end of a 3-ml syringe in RPMI 1640 (Gibco, Gaithersburg, MD). For bone marrow isolation, PBS was injected using a 21G needle to flush cells. Cells were spun and red blood cells were lysed using ammonium-chloride-potassium (ACK) lysis buffer (8.3g ammonium chloride, 1g Potassium Chloride, 250 µl of 0.5M EDTA made to 1000 ml with water PH 7.2-7.4). Cells were then counted using a hemocytometer (Hausser Scientific, Horsham, PA). To only count live cells, trypan blue exclusion was used (Gibco, Gaithersburg, MD). Brains were mechanically homogenized using the dounce method followed by 38% percoll (Sigma, St. Louis, Mo) centrifugation as previously published(*48*). Total cells recovered were counted on the hemocytometer with trypan blue (Gibco, Gaithersburg, MD). Cells were then stained with appropriate antibodies for 30 minutes at 4°C and washed twice with phosphate-buffered saline (PBS, Corning, Corning, NY) and analyzed by flow cytometry (BD, Franklin Lakes, NJ).

### Mouse blood collection and processing

A 1000 unit heparin (Sigma, St. Louis, MO) solution was made in water. This solution was then diluted 1:4 with PBS to generate heparin working solution. Mice were placed in a restrainer and a shallow cut was made over the tail vein using a sterile razor blade. 20-80 µl of blood was collected using a small transfer pipette (Samco Scientific, San Diego, CA, Cat # 231) and immediately placed in the heparin working solution. Heparinized blood was then centrifuged at 400 *g* for 5 minutes and liquid was discarded. Pellet was resuspended in 2 ml of ACK lysis buffer for 3 minutes and quenched with PBS (Corning, Corning, NY) followed by 2 washes. Pellets were then stained with a panel of antibodies for 30 minutes (4°C) followed by washes in PBS before analysis by flow cytometry (BD, Franklin Lakes, NJ).

### Antibodies and Flow cytometry

FITC anti CD8 (BD pharmingen, San Jose, CA, Cat# 553031), FITC anti MHCII (Biolegend, San Diego, CA, Cat# 107605), PE anti CD44 (BD Pharmingen, San Jose, CA, Cat#553134), APC anti CD4 (Biolegend, Cat# 100515), PE-Cy7 anti CD8a (BD, Cat # 552877), PE-Cy7 anti TCRβ (eBioscience San Diego, CA, Cat # 60-5961), Percp anti CD45 (Biolegend, Cat #, 103130), BV785 anti CD8 (Biolegend, Cat # 100750), BV786 anti CD4 (BD Horizon, Cat # 563727), Pacific Blue anti IA/IE (Biolegend, Cat # 107620), BV421 anti CD25 (Biolegend, Cat # 102033) antibodies were used at 1:100 dilution to stain cells isolated from all organs, and at 1:1000 dilution to stain blood cells. Ghost red viability dye (Tonbo, San Diego, CA, Cat # 13-0865-T100) was used at a 1:1000 concentration to stain dead cells. Cells were analyzed using an LSRII flow cytometer (BD, San Jose, CA). Data was analyzed on FlowJo 10.1 (FlowJo LLC, Ashland, OR).

### CFSE labeling and *in vitro* T cell proliferation assay

cells from spleens, inguinal, brachial, and axial lymph nodes of naïve un-manipulated WT C57BL/6 mice were isolated and processed in RPMI 1640 (Gibco) as above. Red blood cells were lysed using ACK buffer and quenched with RPMI (1ml/min/spleen). Cells were then placed in Robosep buffer (Stemcell technologies, Vancouver, BC) at 23-50 million/ml. 1 µl/ml of CFSE (Biolegend, 5mM in DMSO stock) was added at a final concentration of 5 µM. Cells were incubated with CFSE for 10 minutes at room temperature and immediately quenched with complete RPMI (L-glutamine, 10% FBS and 1 % Penicillin/Streptomycin). Cells were washed in complete RPMI twice, counted, and resuspended at 100 million/ml in Robosep buffer for T cell isolation. T cell isolation was performed using EasySep mouse negative T cell isolation kit (Stemcell technologies, Cat # 19851) according to manufacturer’s guidelines. Isolated T cells were counted and resuspended in complete RPMI. Cells were then plated either at 700000 cells/well in a 96-well plate, or at 1.6 million cells/well in a 24-well plate. Anti CD3/CD28 beads (Dynabeads, Gibco, Cat # 11452D) were added at either 19 µl/well for 96-well plates or at 75 µl/ well for 24 well-plates. Total volume was 1 ml in 24-well plates and 250 µl in 96-well plates. To add serum from naïve mice or mice with neurological trauma to these cultures, we added 12.5 µl of serum to 96-well plates or 50 µl of serum in 24-well plates. 72 hours later, T cells were separated from beads using a magnet (Stemcell technologies, Vancouver, BC) and stained for CD4, CD8 and viability dye. CFSE dilution was analyzed by flow cytometry and used as a measure of T cell activation and proliferation.

### Molecular weight analysis of serum

Serum was isolated from naïve mice or mice with a detectable GL261 tumor. Sera was then diluted and passed through a 3kDa filter (Cat # 88512 Pierce ThermoSicentific Waltham, MA) according to manufacturer’s instructions. Top and bottom fractions were collected and separately tested in CFSE assays. Top fragment (>3KDa) was then passed through a 10 kDa (Cat # 88513 Pierce Thermo Scientific Waltham, MA) filter and fractions were tested as before. Top fraction (>10Kda) was then passed through a 30 kDa (Cat # 88502 Pierce Thermo Scientific Waltham, MA) filter and tested as before. Finally, the top fraction was passed through a 100 kDa (Cat # 88503 Pierce Thermo Scientific Waltham, MA) filter and its immunosuppressive ability was tested as before.

### RNA extraction and DNase treatment

Thymi for RNA sequencing were flash frozen in liquid nitrogen immediately upon extraction from the mouse and were kept at −80 °C until the time of isolation. For RNA extraction, frozen thymi were directly placed in 1 ml of Trizol reagent (ambio, Life technologies, Carlsbad, CA) and homogenized using an electric homogenizer (Fisherbrand 150 homogenizer). Once the tissue was completely homogenized, it was incubated at room temperature for 5 minutes. 200 µl of Chloroform (Sigma, St. Louis, MO) was added and samples were vortexed for 15 seconds. Samples were spun at 12000*g* (4°C) for 15 minutes and the top aqueous layer was then collected and mixed with an equal volume of isopropanol (Fisher), vortexed again and spun at 12000*g* (4°C) for 30 minutes to pellet the precipitated RNA. All the liquid was taken off and the pellet was resuspended in 750µl of 75% ethanol. Samples were spun again at 12000*g* for 5 minutes, supernatant was removed, and pellets were dried and resuspended in nuclear-free water. Concentration of RNA was determined using a Nanodrop spectrophotometer. 3 µg of total RNA were treated with DNase I (Invitrogen, DNase1 amplification grade Cat # 18068-015) according to the manufacturer’s instructions. Impurities were eliminated using RNeasy columns (Qiagen, Hilden, Germany, RNeasy Mini kit Cat# 74104) according to the manufacturer’s instructions.

### RNA quality check by methylene blue staining

3 µg total RNA were loaded on a 1% agarose/formaldehyde gel (MOPS buffered) and separated at 80 V for 3.5 hrs. The RNA was transferred onto a positively charged nylon membrane using capillary transfer with 20x SSC overnight and crosslinked by UV (0.125 J/cm2). Membranes were stained with 0.02% methylene blue in Na-acetate buffer and washed with DEPC H_2_O as previously published(*49*).

### Library preparation and RNA sequencing using Illumina TruSeq v2 mRNA Protocol

RNA libraries were prepared using 100 ng of good quality total RNA according to the manufacturer’s instructions for the TruSeq RNA Sample Prep Kit v2 (Illumina, San Diego, CA) employing poly-A mRNA enrichment using oligo dT magnetic beads. The final adapter-modified cDNA fragments were enriched by 12 cycles of PCR using Illumina TruSeq PCR primers. The concentration and size distribution of the completed libraries was determined using a Fragment Analyzer (AATI, Ankeny, IA) and Qubit fluorometry (Invitrogen, Carlsbad, CA). Libraries were sequenced at 30-40 million fragment reads per sample following Illumina’s standard protocol using the Illumina cBot and HiSeq 3000/4000 PE Cluster Kit. The flow cells were sequenced as 100 × 2 paired end reads on an Illumina HiSeq 4000 using HiSeq 3000/4000 sequencing kit and HD 3.4.0.38 collection software. Base-calling was performed using Illumina’s RTA version 2.7.7.

### RNA sequencing analysis

We employed tools available on the Galaxy platform (https://usegalaxy.org) for RNA-Seq analysis. BAM files with unaligned reads were converted to FASTQ files (SAM-to-FASTQ v. 1.126.0)(*49*). The paired-end reads were aligned to the mouse reference genome (mm10; GRCm38.p6; GenBank accession number GCA_000001635.8) using HISAT2 program to generate aligned BAM reads. Aligned BAM files were also visualized with IGV (v. 2.3.98). Using DEXSeq-Count (v. 1.24.0.0), exon read count tables were generated and further analyzed in Microsof Excel 2010. The counts were normalized per million total reads and Log2-transformed. A two-dimensional principal component analysis (PCA) was then performed using Microsoft Excel. To evaluate significant changes in the gene expression patterns, we performed one-way ANOVA for each gene, followed by a two-tailed Student’s t test for each combination of groups. We also calculated fold changes of gene expression compared to the naïve group for a subset of genes involved in thymic function. Pathway enrichment was calculated for genes that were significantly up- or down-regulated at least 2-fold.For this we used the Panther GO-slim pathway analysis tool at www.pantherdb.org. In addition we performed ingenuity pathway analysis to investigate biological function and disease information and associated canonical pathways of interest within our RNA-Seq data (Ingenuity® Systems, www.ingenuity.com).

### Ingenuity pathway analysis (IPA)

The analyzed data from above was uploaded into IPA for further analysis. Canonical pathway analysis was performed using the Ingenuity pathway analysis software (IPA) (Ingenuity® Systems, www.ingenuity.com). Biological functions and disease information within the IPA software were used to investigate the canonical pathways of interest.

### Parabiosis

Two female mice of similar weight and age were co-housed for two weeks prior to surgery to ensure harmonious cohabitation. Anesthesia was induced and maintained with Ketamine/Xylazine (as before). We applied ophthalmic ointment to prevent dry eyes. For analgesia during surgery, we administered Carprofen and Buprenorphine-SR subcutaneously at a dose of 5 mg/kg and 1 mg/kg respectively. We placed the animals on the supine position and shaved the left side of one mouse and the right side of the other mouse from 1 cm above the elbow to 1 cm below the knees. This area was then aseptically prepared with Betadine. During surgery, mice were placed on a heat pad covered by a sterile drape. Animals were then placed on their side, back to back, with adjacent shaved areas facing up. Using a sharp scissor, we performed longitudinal skin incisions to the shaved sides of each animal starting at 0.5 cm above the elbows all the way to 0.5 cm below the knee joints. Following incision, we gently detached the skin from the subcutaneous fascia by lifting the skin up with a pair of curved forceps dissecting the fascia with a second pair to create 0.5 cm of mobilized skin. Next, we began the joining by attaching the left olecranon of one animal to the right olecranon of the other. To facilitate the joining, we bent the elbows of the first and second mouse and passed the needle of a non-absorbable 3-0 silk suture (Ethicon Inc, Somerville, NJ) under the olecranon. We attached joints tightly by a double surgical knot. We next connected the knee joints following the same procedure. Following the attachment of the joints, we connected the skin of the two animals with a continuous absorbable 4-0 vicryl suture (Ethicon Inc, Somerville, NJ). Suturing was started ventrally from knees towards the elbows. Once the ventral skin attachment was completed, we flipped the mice and performed identical suturing on the opposite side going from elbows to the knees until the entire wound was closed with double surgical knots. We verified the continuity of sutures and lack of any openings. We administered 0.5 ml of 0.9% NaCl subcutaneously to each mouse to prevent dehydration. Following recovery, we provided analgesics Carprofen and Buprenorphine SR subcutaneously. Prophylactically, we treated mice with Enrofloxacin (50 mg/kg) in their water bottle for 10 days to prevent bacterial infections. This protocol was adapted from Kamran, et al, as published in JOVE(*50*) and modified according to the IACUC at Mayo Clinic.

### Statistical analysis

GraphPad prism 8.0.1 (La Jolla, CA) was used for figure generation and statistical analysis. Either Mann Whitney U rank sum test or a one way Anova with post hoc analysis or Chi Square survival analysis was used to assess statistical significance. The specific test used is detailed in figure legends. All data are represented as individual data points with mean ± standard deviation (SD).

## Results

### Experimental models of CNS cancer induce sustained thymic involution

CNS cancers are associated with severe immunosuppression (*1, 2, 6*). We sought to determine if this immunosuppression affects the thymus. Specifically, we sought to determine if chronic neurological injuries such as cancers of the CNS induce thymic involution. To test this, we used the GL261 murine glioma model of GBM (Figure 1 A). Additionally, we used intracranial inoculation of B16 melanoma cells as a second model of CNS cancer (Figure 1A). We investigated the effect of brain injury due to CNS cancers on the thymus at the time mice became moribund. GL261 glioma inoculation is incurable and all tumor-bearing mice succumb to death within 50 days (Figure 1 B-survival of tumor-bearing mice is shown). Tumor incidence with GL261 gliomas is between 70-90%. As a result, 10-30% of inoculated mice do not develop tumors and serve as tumor negative controls (tumor ^-^). The thymi of late stage GL261 tumor-bearing mice appeared significantly smaller compared to time-matched/age-matched controls (Figure 1A). In a larger cohort, we determined that thymic size as measured by weight and cellularity was significantly reduced in GL261 tumor-bearing mice when compared to tumor ^-^ controls (Figure 1 C-D). Similarly, GL261 implanted mice had smaller thymi (as measured by weight and cellularity) when compared to naïve unmanipulated mice (Figure 1 E-F). Subcutaneous implantation of GL261 cells did not induce measurable thymic involution suggesting that unlike CNS tumors, peripheral cancers may not disrupt thymic homeostasis in a similar manner (Figure 1 G). Next we sought to determine whether the extent of thymic involution is dependent on the magnitude of brain injury. To determine whether thymic involution was related to the degree of brain injury, we quantified thymic cellularity in the GL261 glioma model. We took advantage of the heterogeneity in GL261 glioma growth in our experiments as mice bearing luciferase-expressing GL261 tumors have varying tumor burdens at the onset of morbidity. This variation can be measured using bioluminescence imaging. Based on the luminescence intensity, mice were grouped into low, medium, and high tumor burden cohorts. Thymic cellularity was then correlated with tumor burden in these mice. As shown in Figure 1H, we found an inverse correlation between thymic cellularity and tumor burden.

**Figure 1:**
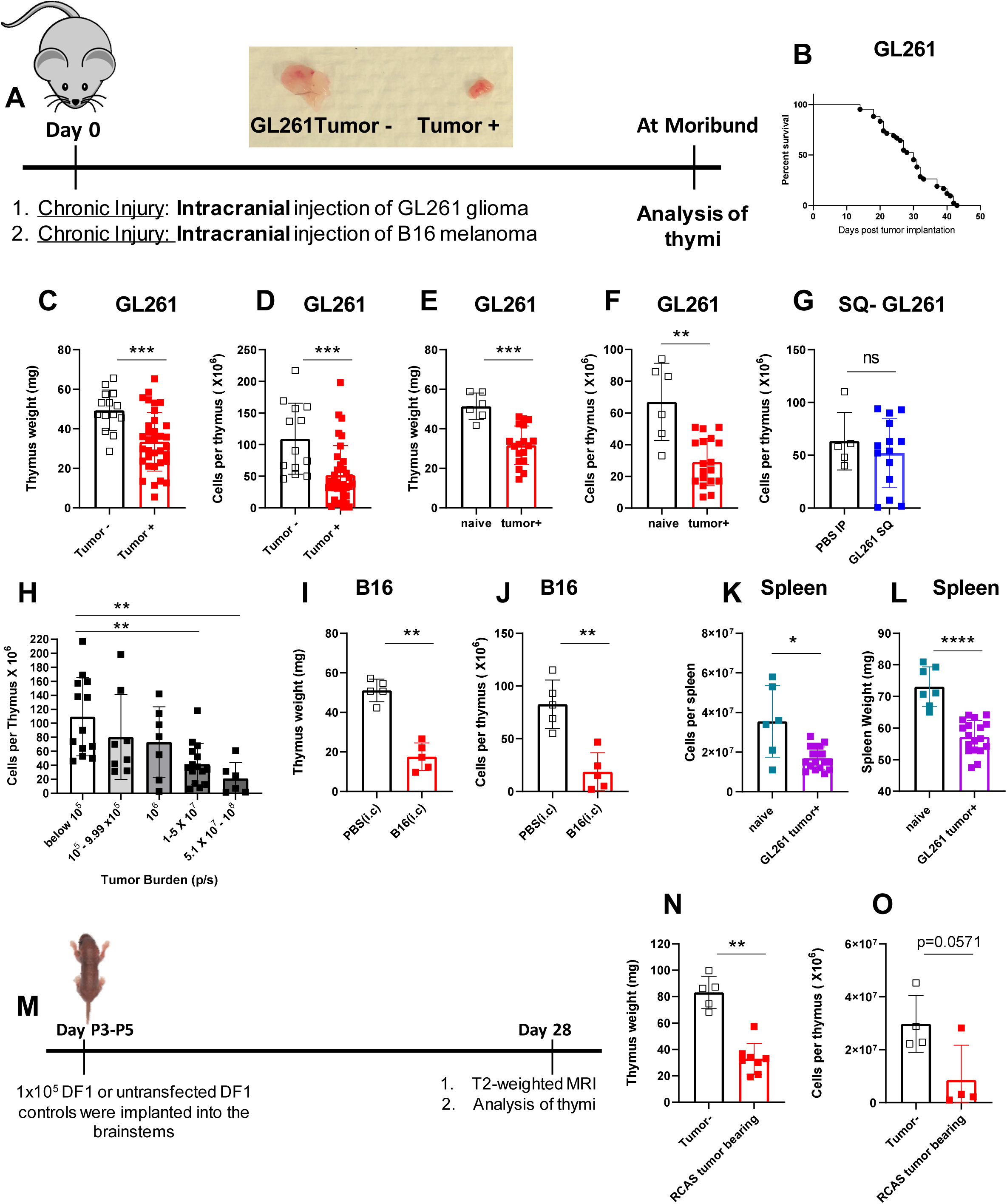
CNS cancer is a chronic neurological insult that results in sustained thymic involution: Experimental design is shown in (A). Mice are implanted with 60,000 luciferase-bearing cells or 10,000 B16-F1 melanoma cells in the frontal lobe of the brain 3 mm deep. Gross analysis of thymi taken from a GL261 tumor bearing mouse compared to a tumor negative mouse (A)Thymi are analyzed at the time mice become moribund. Survival of GL261 bearing mice is shown in (B). Tumor take is 70-90% rendering the remaining 10-30% tumor negative. Thymic weight (C) and cellularity (D) is compared between GL261 tumor bearing and tumor negative mice at the time glioma-bearing mice become moribund. Tumor negative and tumor-bearing mice are age-matched. Similarly GL261 bearing mice have significantly smaller thymi as measured by thymic weight and cellularity compared to naïve unmanipulated mice (E-F). Subcutaneous implantation of GL261-Luc cells did not result in thymic involution (G). Thymic cellularity was inversely correlated with tumor burden in GL261-Luc bearing mice (H). Mice were binned based on bioluminescence imaging at the time of euthanasia and thymic cellularity was plotted against tumor burden (as measure by bioluminescence signals). Thymic weight (I) and cellularity (J) are compared between PBS injected and B16 melanoma implanted mice on day 11 when B16 implanted mice became moribund. Similar to the thymus, spleens of Glioma bearing mice (GL261 implanted) are significantly reduced compared to naïve unmanipulated controls (K shows weight –L is cellularity). Experimental design for implantation and analysis of RCAS spontaneous gliomas is shown (M). Thymic weight (N) and cellularity (O) in tumor negative vs. RCAS tumor-bearing is shown. N=4-8 for RCAS experiments. n=13-34 in GL261 experiment pooled from 3 to 4 independent experiments. n=5 in B16 experiment. All B16 implanted mice developed detectable tumors. Data are shown as individual mice with mean. Error bars represent standard deviation. Mann-Whitney test was used to assess statistical significance. ns p ≥ 0.05, * p= 0.01 to 0.05, ** p=0.001 to 0.01, ***p=0.0001 to 0.001, **** p< 0.0001.

We next sought to examine whether our findings were glioma-specific. Similar to GL261 gliomas, implantation of B16 melanoma in the brain induced a significant thymic involution compared to sham injected mice as evident by reductions in both thymic weight and cellularity (Figure 1 I-J). In addition, the thymus was not the only immune organ affected by brain tumors. We also found that spleens of tumor-bearing mice (GL261) were reduced in size and cellularity compared to naïve unmanipulated controls (Figure 1 K-L). Finally, we tested the extent of thymic involution in the RCAS based spontaneous murine glioma model of diffuse intrinsic pontine glioma (DIPG) as described in the methods section (Figure 1 M). In this model, 3-5 day old mice were intracranially injected with cells that can induce spontaneous glioma growth in the brain stem. One month later, mice were analyzed by T2-weighted MRI to determine the presence or absence of a tumor (Figure 1 M). Thymic involution was observed in mice that spontaneously developed gliomas in the brain stem compared to those that did not (Figure 1 N-O). Taken together, our results demonstrate that three distinct experimental CNS cancers, which grow in different brain regions, cause similar thymic involution. These data also show that GL261 glioma size correlates with the degree of thymic involution.

### Brain tumors induce changes within the thymus that affect developing T cells at distinct stages of development

Given that GBM growth in the brain induced severe thymic involution, we next sought to determine cellular changes within the affected thymi. We first compared age-matched tumor ^-^and naïve thymi and did not find any differences between these groups in our experiments (data not shown). Therefore, both tumor ^-^and naïve mice were used as controls interchangeably. To analyze T cell development within the thymus, we gated on live, CD45^+^ hematopoietic cells and used CD4 and CD8 markers to determine double positive, DP, (CD4^+^ CD8^+^), single positive, SP4, (CD4^+^ CD8^-^), SP8 (CD4^-^ CD8^+^), and double negative, DN, (CD4^-^ CD8^-^) populations (Figure 2 A). Once DN populations were identified, we used CD44 and CD25 markers (within the cells in the DN gate) to distinguish DN1 (CD44^+^ CD25^-^), DN2 (CD44^+^ CD25^+^), DN3 (CD44^-^ CD25^+^), and DN4 (CD44^-^CD25^-^) populations of developing thymocytes (Figure 2 B). Using these gating strategies, we assessed the stage at which T cell development is affected by a growing glioma in the brain. Frequencies of DN1 cells were significantly increased in GL261 tumor-bearing mice compared to controls (Figure 2 C). Interestingly, this increase in frequencies of DN 1 cells translated into normalization of absolute numbers of DN1 cells between control and atrophied thymi (Figure 2 C). Frequencies of DN2 cells trended towards an increase though this difference did not reach statistical significance (Figure 2 D). Similar to DN1 cells, we observed analogous numbers of DN2 cells in both control and atrophied thymi (Figure 2 D). Interestingly, both frequencies and absolute counts of DN3 cells were significantly reduced in tumor-bearing mice compared to controls (Figure 2 E) suggesting T cell development is blocked at the DN2-DN3 transitional stage. This reduction in DN3 cells translated into a significant reduction in frequencies and numbers of DN4 cells as well as DP cells (Figure 2 F and G). Intriguingly, frequencies of single positive CD4 T cells were significantly increased in glioma-bearing mice compared to controls (Figure 2 H). This significant increase in frequencies of SP CD4 T cells translated into the equalization of absolute numbers between control and atrophied thymi (Figure 2 H). Frequencies of SP CD8 T cells were also increased in glioma-bearing mice compared to controls (Figure 2 I). However, this increase in frequencies of SP CD8 T cells was not sufficient to completely compensate for the loss of cells and hence absolute numbers of SP CD8 T cells in glioma-bearing mice remained reduced when compared to controls (Figure 2 I).

**Figure 2:**
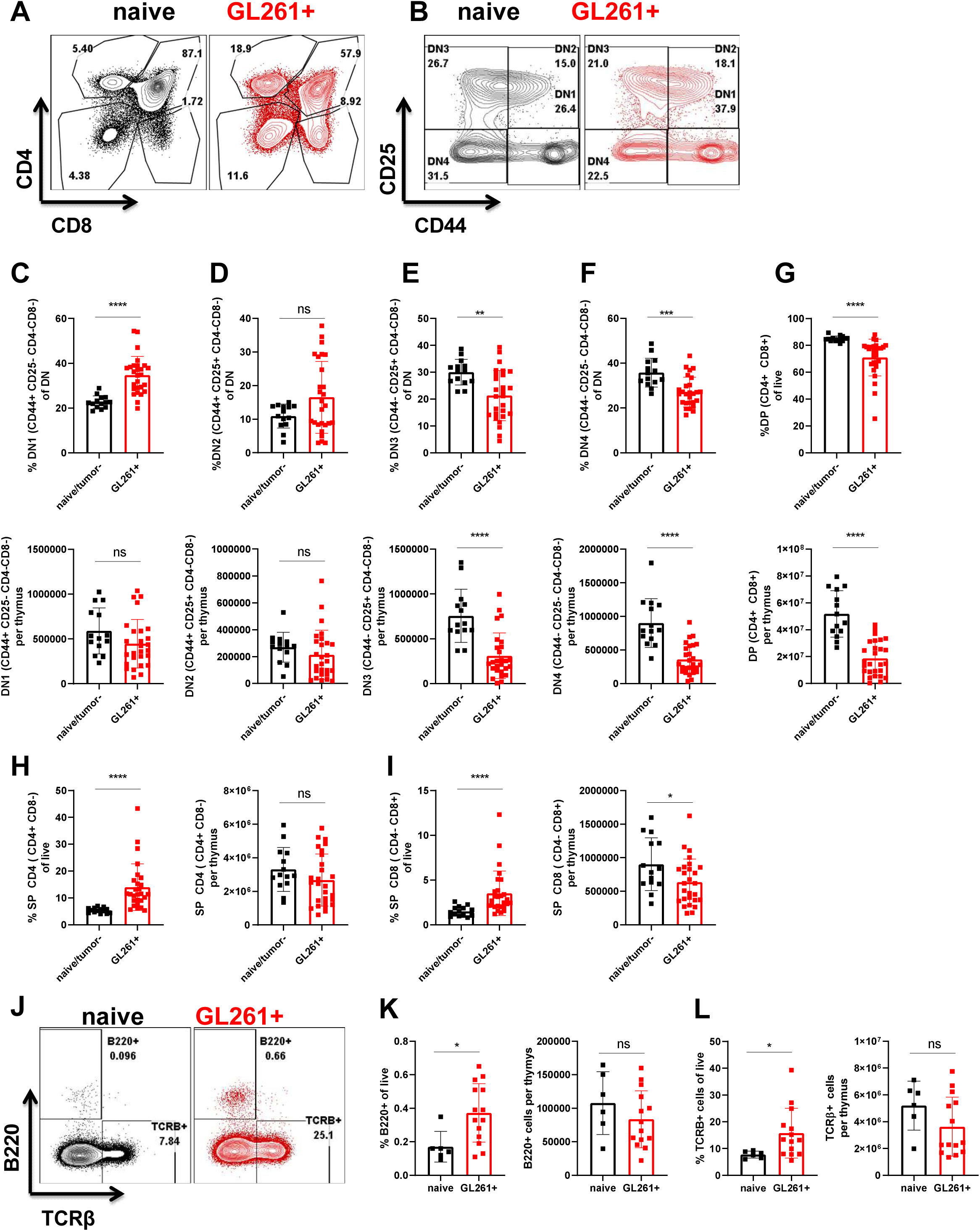
T cell development programs are halted at the DN3 stage of development in glioma-bearing mice compared to controls: Gating strategy is shown using representative naïve and glioma-bearing mice in A and B. Following gating on singlets, live, and CD45+ cells, we used CD4 and CD8 parameters to determine DN (double negative), DP (double positive), SP4 (single positive CD4), and SP8 (single positive CD8 T cells) (A). We then focused on the DN gate to define DN1-4 populations (B). DN1 is defined as CD44+ CD25-, DN2 is defined as CD44+ CD25+, DN3 is defined as CD44-CD25+, and DN4 is defined as CD44-CD25-populations within the DN gate (B). Frequencies (top) and numbers (bottom) of DN1-DN4 and DP cells are quantified between naïve or tumor- and glioma bearing mice (C-G). N=14-27. Data are shown as individual mice with mean. Data are pooled from 2 independent experiments. This experiment was repeated a total of 5 times and similar results were obtained. Error bars represent standard deviation. Mann-Whitney test was used to assess statistical significance. Ns p ≥ 0.05, * p= 0.01 to 0.05, ** p=0.001 to 0.01, ***p=0.0001 to 0.001, **** p< 0.0001. Post selection TCRβ+ cells and B220+ B cells within the thymus are increased in glioma-bearing mice compared to controls (J). Frequencies and numbers of thymic B cells are quantified in (K). Frequencies and numbers of TCRβ+ cells in the thymus are quantified in (L). N=6-14. Data are shown as individual mice with mean. Error bars represent standard deviation. Mann-Whitney test was used to assess statistical significance. Ns p ≥ 0.05, * p= 0.01 to 0.05, ** p=0.001 to 0.01, ***p=0.0001 to 0.001, **** p< 0.0001.

B cell developmental programming is actively inhibited in the normal thymus to maximize generation of functional T cells (*51–53*). Hence, we evaluated frequencies and numbers of thymic B cells and SP T cells that have successfully rearranged their T cell receptor using flow cytometry (Figure 2 J). Interestingly, we found a significant increase in frequencies of both B cells and TCRβ ^+^ T cells in the atrophied thymi obtained from glioma-bearing mice compared to controls (Figure 2 K-L). This significant increase in frequencies of B cells and TCRβ ^+^ T cells resulted in normalization of absolute counts of these populations between control and atrophied thymi (Figure 2 K-L). It is worth mentioning here that thymic involution due to stress responses, peripheral infections, and aging is thought to mainly affect DP cells (in cases of stress responses) and/or epithelial cells (in aging)(*54*). While a reduction in DP cells is associated with stress induced thymic involution (*54–56*), a DN2-DN3 block with enhancement of single positive cells and B cells is not consistent with stress induced thymic involution. Thus our data indicate that experimental GBM induces thymic involution with a unique cellular signature. This signature is associated with inhibition of T cell development at the DN3 stage, accumulation of SP cells, and enhancement of B cell development in the thymus.

### Thymi of mice with experimental GBM present with unique gene transcription patterns compared to naïve mice

We observed thymic involution in GL261 glioma-bearing mice that was associated with a DN3 block in T cell development, retention of SP T cells, and enhancement of B cell development at the cellular level. We next asked whether thymic involution in glioma-bearing mice was associated with distinct transcriptional programs in the thymus. We also aimed to determine whether thymic involution due to CNS cancer resembles those changes observed in stress or aging mediated pathways. We employed RNA sequencing as an unbiased method of analysis that can identify upstream pathways and/or molecules that regulate thymic involution. We used next generation RNA sequencing of the transcriptome followed by gene onthology (GO) and Ingenuity pathway analysis (IPA) to determine if thymi of glioma-bearing mice were transcriptionally distinct from thymi of sham control and naïve unmanipulated mice. To accomplish this, we inoculated C57BL/6 mice with GL261 glioma cells or injected PBS as sham control (Figure 3 A). At the time a tumor-bearing mouse became moribund, we euthanized that mouse as well as one sham and one naïve time-matched/age-matched controls and isolated bulk thymic RNA. Three GL261 tumor-bearing, three PBS sham, and three naïve thymi that had the best quality of RNA were selected for RNA-sequencing (Figure S1 A). Quality of RNA was assessed by degradation status of the RNA (Figure S1 A). We used principle component analysis (PCA) to determine if the three groups were distinct from each other. Both 2 dimensional and 3 dimensional PCA analysis placed thymi of glioma-bearing mice separate from both PBS and naïve groups (Figure 3 B and data not shown) suggesting gene transcriptional programs are significantly different in thymi of mice with on-going brain insults. It is worth mentioning that at this time (between days 25-42 post intracranial injection), we did not observe separation between PBS and naïve groups (Figure 3 B and Figure S1 C).

**Figure 3:**
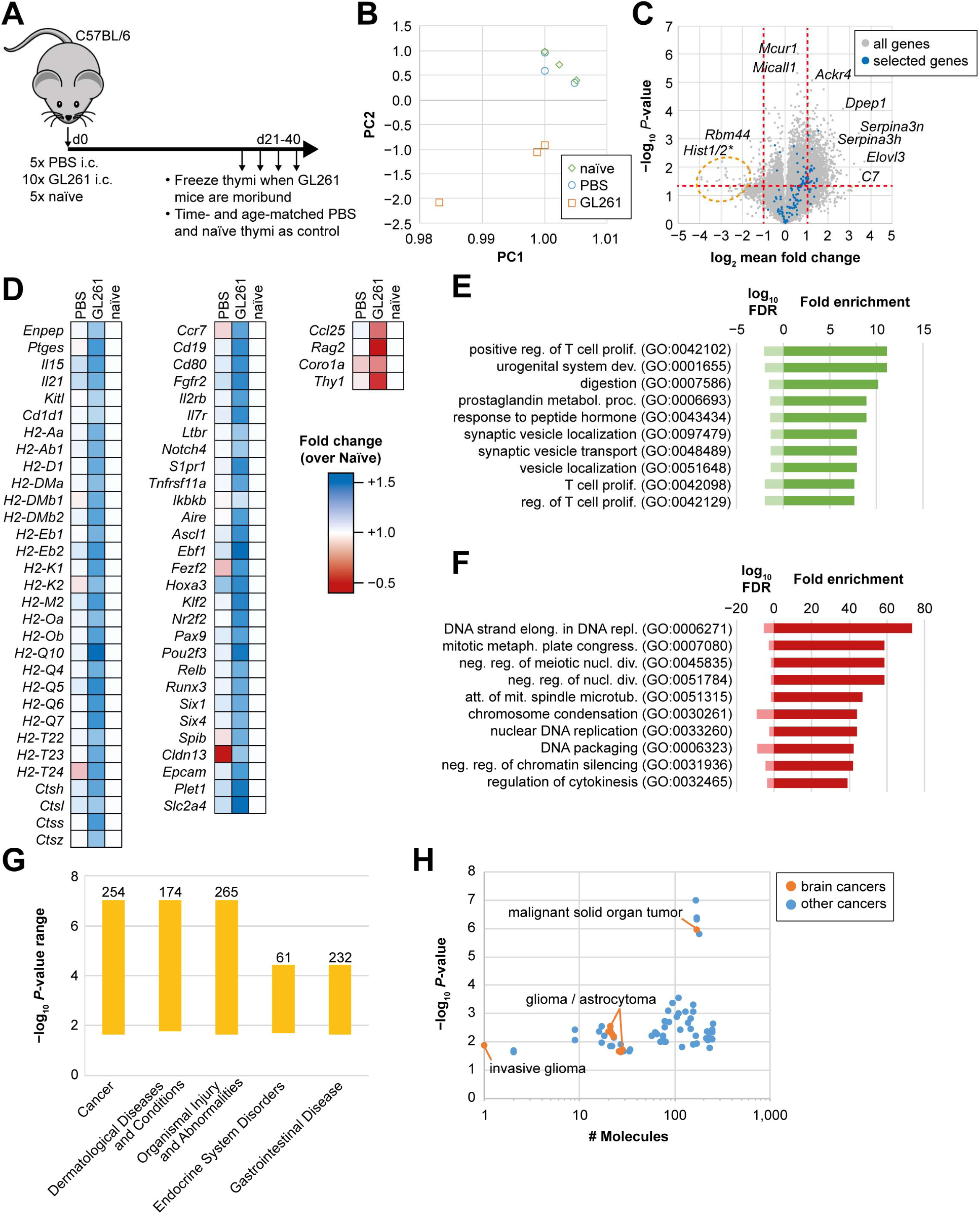
Thymic epithelial cell (TEC) signatures are enhanced in glioma bearing thymi while T cell commitment programs are blocked at DN2-DN3 stage of development. Experimental design for RNA-sequencing experiment is shown (A). Three naïve, three PBS injected (time and age matched with glioma-bearing mice) and three GL261 implanted mice were chosen from 5 or 10 mice based on quality of RNA extracted from thymus. RNA from these 9 mice was used to perform next generation sequencing. Principle component analysis of data sets demonstrates distinct transcriptional programs between GL261 and sham or naïve controls (B). This is while naïve and sham injected mice (that have recovered from injury by the time of harvest) cluster closely. Expression levels of genes differentially regulated between GL261-bearing thymi and naïve thymi are shown in (C). Grey dots represent all genes. Blue dots represent gene sets used for analysis of genes specifically involved in epithelial cell development, maturation, and function, T cell lineage commitment, cytokine signaling and other factors reported in the literature to influence thymic involution in several models. Highly regulated genes (up or down) are shown in figure overlaying the dots. (D) Blue genes from C are further statistically analyzed and shown in heat map. This data is normalized to levels of expression in naïve thymi. White box represents a value of 1 relative to naïve. Blue represents upregulation and red represents downregulation. For statistical analysis, we use a one-way ANOVA for every gene and calculated the p-value based on significance and change in expression. Results from gene enrichment analysis and pathway analysis are shown in E for significantly upregulated pathways and in E for significantly downregulated pathways. Panther Slim-GO website was used for this analysis. (E) Shows analysis of all 1404 genes that were significantly upregulated at least 2 folds. The dark bars show fold enrichment of the pathway members in this group (calculated as (#identified hits / #analyzed genes) / (# genes in the pathway / # total mouse genes). The light bar shows the log10 value of the false discovery rate. The analysis is restricted to the diagram to the top 10 hits. Pathways are ordered by fold enrichment. Most involved pathways upregulated are related to T cell development and proliferation, and hormone signaling. (F) Here we demonstrate an analogous analysis of significantly downregulated genes. The involved pathways are mostly DNA replication, cell cycle and cell division-related. Ingenuity pathway analysis is used to unbiasedly determine the top diseases associated with our data set. We found cancer to be the top diseases associated with our dataset with 254 molecules known to regulated cancer being enriched in our data set (G). Cancers that are likely regulated by the enriched in our data set are further subdivided in (H) with glioma and brain tumors being represented amongst the IPA hits.

We found a significantly altered gene expression pattern in thymi from GL261 bearing mice compared to naïve mice (Figure 3 C). The changes in thymi of glioma-bearing mice were also highly consistent when compared to PBS-injected mice (Figure S1 B), while no significant differences were observed between the two control groups (Figure S1 C). Genes with highest levels of significant up or down regulation are labeled in the plot (Figure 3 C). Interestingly, genes highly upregulated include *Serpina3n* and genes related to the complement pathway. These genes have not been reported to be regulated by stress induced or age mediated thymic involution (*54, 55, 57-61*). Instead, these genes were implicated mainly in pathology of neurological diseases including pain, peripheral nerve damage, and recently in Alzheimer’s disease (*62–67*).

### Thymic epithelial cell (TEC) signatures are enhanced in glioma bearing thymi while T cell commitment program signatures are blocked at the DN2-DN3 stage

#### Epithelial cell gene expression is normal in atrophied thymi of GBM-bearing mice

We next investigated the expression changes of a set of genes known to be important in thymic T cell development, epithelial cell development, epithelial cell maturation and function, and cytokine signaling (Figure 3 D: also indicated as blue dots in Figure 3 C). Levels of expression were normalized to naïve mice (white boxes represent values of 1). Genes significantly upregulated in mice intracranially injected with GL261 or PBS are represented in heat maps with blue representing a significant upregulation and red representing a significant down regulation (Figure 3 D and Figure S1 D). We did not find a signature associated with epithelial cell death and degeneration (Figure 3 D and S1 D) as reported in thymic involution due to aging (*54, 61, 68*). Instead, we observed upregulation of genes related to TEC progenitors and lineage commitment (*Plet1*, *Claudin3*, *Claudin4*, *Relb*), TEC proliferation and maturation (*Epcam*, MHCII genes, *Cd80*), and several genes important for developmental programs of epithelial cells including *Foxn1* and most genes regulated by *Foxn1* including *Cxcl12*, and *Cd83* (Figure 3 D and S 1 D). Interestingly, some *Foxn1* regulated genes such as *Dll4*, *Ccl25*, and *Psmb11* were downregulated in thymi of GL261 tumor-bearing mice compared to controls (Figure 3 D). *Ccl25* and *Dll4* are mainly important in cTEC interaction with DN thymocytes (*52*). The importance of this downregulation may aid in explaining the defects seen in developing T cells rather than epithelial cells, which will be discussed below.

Mature mTEC genes were also upregulated in thymi of GL261 tumor-bearing mice compared to naïve controls. These genes included *Aire*, *Fezf2*, MHCII genes, and keratins (Figure 3 D and S1 D). *Pax9*, *Six1*, and *Six4* have also been implicated in cTEC development/function and were also significantly increased in thymi of GL261 tumor-bearing mice compared to controls (Figure 3 D and S2 D). Interestingly, homeostatic cytokines and their receptors including *Il15*, *Il21*, *Il7r*, *Il2rb*, *Il7*, and *Il2ra* were all upregulated in thymi of GL261 tumor-bearing mice compared to controls (Figure 3 D and S1 D). Higher expression of these cytokines can indicate availability of fewer T cells to use these homeostatic cues and suggest that involution is likely not due to limited levels of these cytokines.

#### Genes involved in regulation of T cell development and proliferation are reduced in atrophied thymi compared to controls in a manner that supports changes in the cellular compartment

Because RNA-seq was performed on bulk thymi and our data indicated an enhancement of epithelial cell signatures and increased levels of homeostatic cytokines in thymi of GL261 tumor-bearing mice compared to controls, we hypothesized that the involution is due to T cell intrinsic factors and not due to epithelial cell dysfunction as observed in aging models. We next evaluated genes involved in different stages of T cell development. We found that *Rag1* and *Rag2* genes, *TCF3*, *TCF7*, *Gata3*, and *BCl11b*, and *Notch 1* were downregulated in thymi of GL261 tumor-bearing mice compared to controls (Figure 3 D and S1 D). Downregulation of these genes can be consistent with a block between DN2-DN3 transition (Figure 2 and (*52, 53, 69, 70*)). Interestingly, another significantly downregulated gene in thymi of GL261-bearing mice compared to controls was *Coro1a* (Figure 3 D). Coro1a is expressed in hematopoietic cells and regulates T cell exit out of the thymus through modulating sensitivity to S1P (*71*). Coro1a deficiency is associated with accumulation of single positive T cells in the thymus and peripheral lymphopenia (*71*). Downregulation of *Coro1a* in thymi of glioma-bearing mice may explain the accumulation of SP T cells in the atrophied thymi compared to controls (Figure 2 H-I).

#### Gene expression during experimental GBM induced thymic involution is not consistent with a stress response

We next sought to determine biological pathways with enriched genes from our data set. Overall pathways highly upregulated and downregulated according to the level of enrichment in our dataset are shown in Figure 3 E and F respectively. Interestingly, genes enriched in pathways associated with T cell proliferation are highly upregulated suggesting subsets within the thymus are striving to proliferate and maintain cellularity. Genes enriched in pathways that were significantly downregulated included DNA replication and elongation suggesting a possible block in proliferation for some subsets. This signature is inconsistent with published data on pure stress induced proliferation where cell death pathways are involved and highly regulated (*54, 59, 72*). Consistent with this, we evaluated top pathways regulated in our data set as predicted by IPA (Figure S1 F). One of the top 15 regulated pathways was the acute phase response pathway (Figure S1 E-F). In fact, genes associated with glucocorticoid receptor signaling, which is part of the acute response pathway, were not significantly enriched in our data set (Figure S1 E). Instead, genes enriched in this pathway included those encoding *Serpina3*, *Serpina1*, *Il1* signaling, haptoglobin, and fibrinogen (Figure S1 E). Also confirming this, *Nr3c1*, which is the gene encoding for glucocorticoid receptor, was expressed at equivalent levels between glioma-bearing thymi and controls (Figure S1 D).

Finally, we used IPA to determine the likelihood that our data set is associated with particular diseases. Interestingly, analysis of our thymic bulk RNA-seq data from mice with brain gliomas indicated cancer to be the top disease associated with our data set (Figure 3 G). In fact about 254 molecules that are known to regulate cancer development and prognosis were significantly enriched in our thymic RNA data set (Figure 3 G).Interestingly, amongst cancers being linked to our data sets, those affecting the CNS including gliomas, and astrocytomas were highly enriched (Figure 3 H). This is interesting in that bulk RNA-seq of the thymus (an immune organ) can unbiasedly identify similarities between pathways enriched in our dataset and those involved in CNS cancers. This data suggest not only that cancer is a systemic disease, but also that a primary immune organ (the thymus) can be affected by a brain tumor in a manner that is similar to the pathways involved in the cancer itself.

### Mice harboring GL261 gliomas exhibit peripheral immune suppression consistent with GBM patients

We have demonstrated that chronic neurological insults including GBM cause thymic (and splenic) involution. We next investigated if thymic involution occurs concurrently with peripheral immunosuppression affecting other immune organs. Peripheral immunosuppression is observed in patients with GBM and can undermine success of treatments including immunotherapies in these patients. GBM patients often present with low CD4 T cell counts and reduced expression of MHC class II molecule on their peripheral blood monocytes (*2, 6*). Additionally, increased T cell sequestration in the bone marrow and loss of spleen volume was recently reported in patients with GBM (*2*). Because GBM patients have this distinct peripheral immunosuppression phenotype, we further assessed the peripheral immune system in the GL261 model of glioma.

In mice inoculated with GL261 gliomas, we observed a continuous decline in CD4 and CD8 T cells in peripheral blood (Figure 4 A, C-D and S2 B-C). By bleeding the same tumor-bearing mouse weekly (and naïve mice as controls), we were able to observe an overall decline in CD4 T cells as glioma-bearing mice become increasingly moribund (Figure 4 A and Figure S2 A showing two mice; first died on day 14 and the second died on day 21 post tumor inoculation). The drop in frequencies of CD4 and CD8 T cells in mice implanted with GL261 glioma cells translated into reduced T cell counts as total numbers of cells per 100 microliters of blood remained comparable between control and GL261-bearing mice (Figure 4 G-H and S2 D). Frequencies and numbers of B cells in blood trended towards a decline as well, although this difference did not reach statistical significance (Figure S2 E).

**Figure 4:**
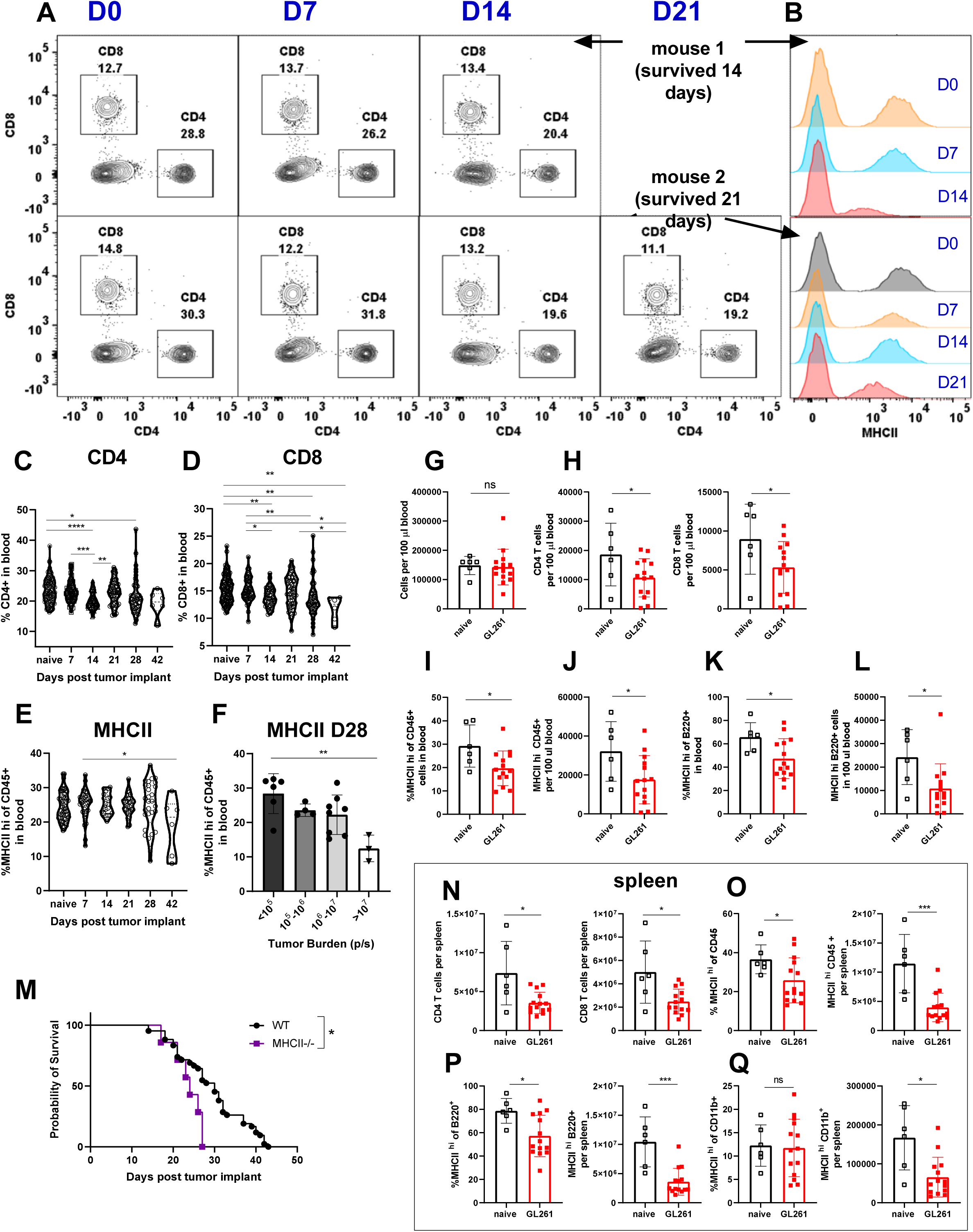
Mice harboring GL261 gliomas exhibit peripheral immune suppression which is consistent with GBM patients. CD4 and CD8 T cell levels are tracked through weekly bleeding in individual mice (A). MHCII expression on all CD45^+^ cells in blood is tracked weekly in individual mice (B). Two representative mice are shown. Frequencies of CD4 (C) and CD8 (D) and MHCII expression on CD45^+^ cells (E) in blood are tracked across GL261 implanted mice through weekly bleeding (n=6-124). Data is pooled from 4-5 independent experiments. Variability in MHCII expression on day 28 is plotted as a function of tumor burden as measured by bioluminescence intensity (I) (n=3-8) data is pooled from 2 independent experiments. Total cells per 100 µl of blood does not change (G) between naïve and glioma bearing. Reduction in frequency of CD4 and CD8 T cells translates into a reduction in total counts of CD4 and CD8 T cells in blood (H). Frequency (I) and number (J) of CD45+ hematopoietic cells in blood with high expression of MHCII is quantified. Similarly, frequency (K) and numbers of (L) B cells with high levels of MHCII expression in blood are decreased in glioma-bearing mice compared to controls. Survival of WT (n=42) and MHCII^-/-^ (n=7) mice implanted with GL261 cells are compared (M). Numbers of CD4, and CD8 T cells (N), and frequencies and numbers of MHCII hi CD45+ (O), MHCII hi B cells (P), and counts of MHCII hi CD11b+ cells (Q) are compared between glioma-bearing and naïve mice in the spleen. For T cell gating: we gated on singlets, live, CD45+, TCRB+ B220-, CD4+ or CD8+ cells. For B cell gating, we used singlets, live, CD45+, TCRB-B220+ cells. For CD11b gating, we used singlets, live, CD45+, TCRB-B220-, CD11b+ cells. Data are shown as individual mice with mean. Error bars represent standard deviation. One way Anova with Tukey’s multiple comparisons test was used to assess statistical significance for C-F. Mann-Whitney-U test was used to assess statistical significance in G-Q. For survival data Log-rank (Mantel-Cox) test was used (MST WT =30 days vs. 24 days for MHCII^-/-^). Ns p ≥ 0.05, * p= 0.01 to 0.05, ** p=0.001 to 0.01, ***p=0.0001 to 0.001, **** p< 0.0001.

We next evaluated MHC class II expression in GL261 glioma-harboring mice. Similar to T cell counts, a drop in MHCII expression and total counts of MHCII positive cells was apparent in all CD45^+^ hematopoietic cells in blood (Figure 4 B, E and J-K). Importantly, this drop in MHCII expression correlated with tumor burden on day 28 as mice with the highest tumor burden had the lowest levels of MHCII expression on their CD45^+^ blood hematopoietic cells (Figure 4 F). This decline of MHCII was also apparent when separately analyzing B cells in the blood (Figure 4 L). MHCII^-/-^ mice had higher tumor incidence post implantation with GL261 cells (data not shown) and lower survival compared to WT counterparts (Figure 4 M) suggesting MHCII downregulation may be a key mechanism of immunosuppression in GBM. Next, we sought to address whether a T cell drop occurs in the spleen as well in glioma-bearing mice compared to controls. Similar to blood, we found a reduction in total numbers of CD45^+^ cells, CD4 T cells, CD8 T cells, and B cells while the frequencies of these populations did not change drastically (Figure 4 O-P, and S2 G-H). This is consistent with the reported literature in a different model of CNS cancer (*2*). Additionally, we observed reduced counts of MHCII^+^ CD45^+^, MHCII^+^ B cells and MHCII^+^ CD11b^+^ monocytes and macrophages (Figure 4 O-Q). We observed a reduction in frequency of MHCII expressing CD45^+^ cells, and B cells, but not CD11b^+^ cells (Figure 4 O-Q).

Finally, we evaluated the bone marrow of glioma-bearing mice for evidence of T cell sequestration as reported in GBM patients and in a different mouse model of glioma (*2*). As expected, we observed a significant, but modest, increase in frequency of T cells within the BM of glioma-bearing mice compared to controls (Figure 5 A-B). A similar trend was observed in total numbers of bone marrow T cells though this difference did not reach statistical significance (Figure 5 B). We did not observe a significant increase or trend in total cellularity of bone marrow suggesting this increase was due to a preferential increase in T cell compartment which is consistent with the notion that T cells are actively sequestered in the bone marrow during neurological insults (Figure S2 I). In accordance with this observation, numbers or frequencies of other cell populations including CD11b^+^ (gated on non B and T cells) and B220^+^ cells did not change between naïve and glioma-bearing mice validating a T cell specific increase in the bone marrow during neurological injuries (Figure S2 J-K). Therefore, GL261-harboring mice exhibit features of peripheral immune suppression similar to GBM patients.

**Figure 5:**
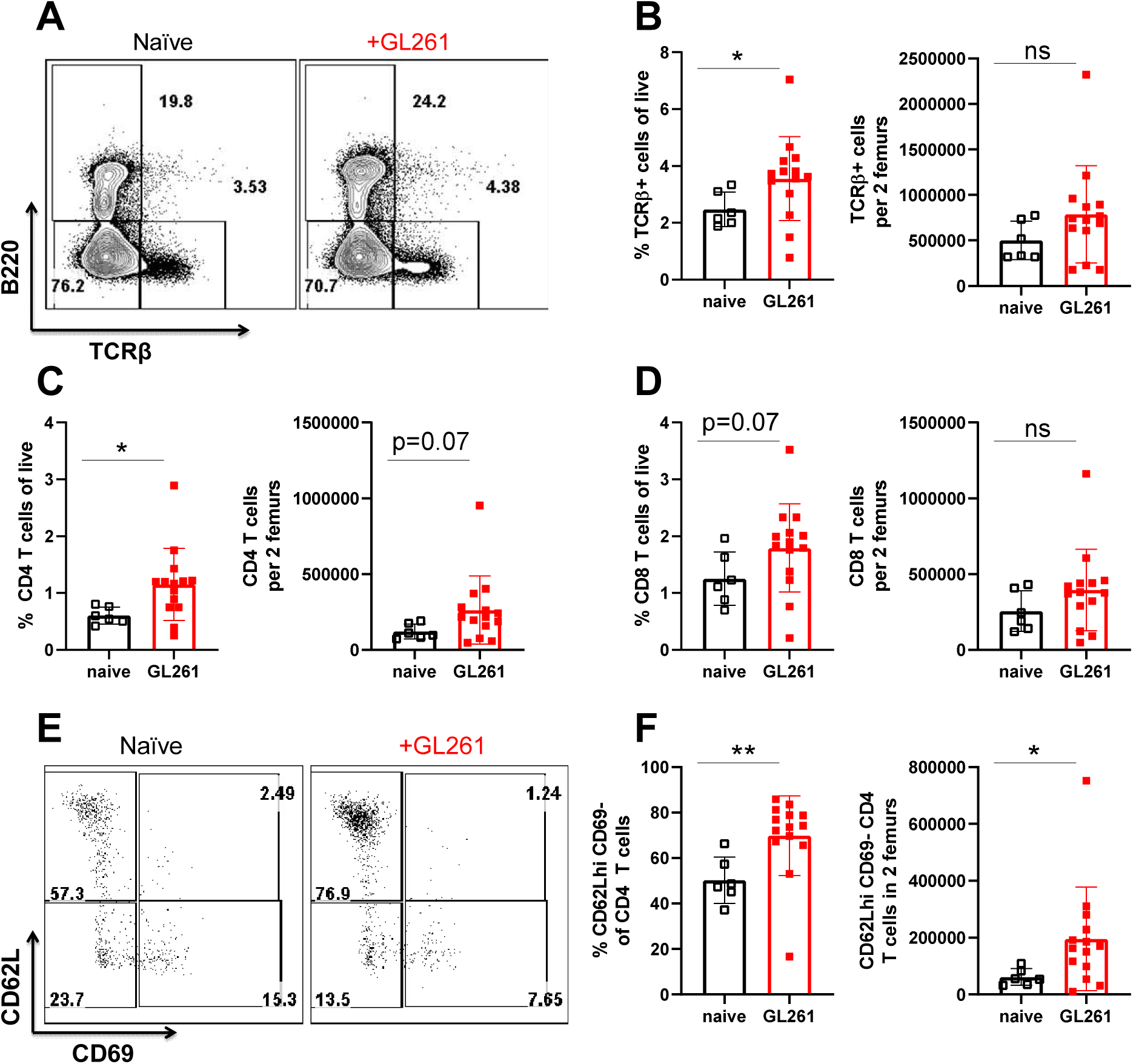
CD4 T cells with a naïve phenotype preferentially accumulate in the bone marrow of glioma-bearing mice A modest number of T cells accumulate in the bone marrow of glioma-bearing mice when compared to naïve controls. (A). Frequency (left) and absolute numbers of T cells (right) in the bone marrow of GL261-glioma-beaing and naïve controls are shown in (B). The increase in the bone marrow T cells is mainly due to an increase in frequencies and numbers of CD4 T cells (C). The increase in bone marrow resident CD8 T cells in glioma-bearing mice does not reach statistical significance (D). Representative flow plot of CD4 T cells within the bone marrow indicates CD4 T cells with CD62L hi CD69-phenotype to accumulate in glioma-bearing mice compared to controls (E). Frequencies and absolute counts of CD62L hi CD69-CD4 T cells are quantified in naïve and glioma-bearing mice (F). GL261+ group represents mice that were intracranially implanted with the glioma cell lines and were verified to be tumor-bearing by bioluminescence. Data presented is at the time GL261+ mice became moribund. Naïve controls are sex/age matched. Two femurs from each animal were collected. For gating, we used a gating strategy that included gates for single cells, live cells, CD45+ hematopoietic cells, TCRβ+ cells, before gating for CD4 and/or CD8 T cells. Within the CD4 gated population, we then used CD62L and CD69 to further investigate the phenotype of bone marrow resident cells. Data is shown from 6 naïve controls and 14 GL261–glioma-bearing mice. Individual data points are shown. Error bars represent standard deviation. Mann-Whitney test was used to assess statistical significance. Ns p ≥ 0.05, * p= 0.01 to 0.05, ** p=0.001 to 0.01, ***p=0.0001 to 0.001, **** p< 0.0001.

### CD4 T cells with a naïve phenotype preferentially accumulate in the bone marrow of glioma-bearing mice

Thus far we recapitulated major hallmarks of glioma-induced immunosuppression in the GL261 model. One such hallmark was accumulation of T cell in the bone marrow of glioma-bearing mice compared to naïve controls. We next sought to determine the specific phenotype of sequestered T cells within the bone marrow during glioma growth. We therefore evaluated whether an analogous increase was observed in CD4 and CD8 compartments within the bone marrow of glioma-bearing mice compared to controls. We found that although both bone marrow resident CD4 and CD8 T cell frequencies and numbers trended towards an increase in glioma-bearing mice, the effect on CD4 T cells appeared more pronounced and reached statistical significance (Figure 5 C-D). We next investigated the specific population of CD4 T cells that accumulate in the bone marrow of glioma-bearing mice. We determined that the naïve bone marrow harbors populations of CD62L^+^ CD69^-^ CD4 T cells at baseline (Figure 5 E-F). Interestingly, this same population is expanded in glioma-bearing mice when compared to naïve controls (Figure 5 E-F).

Together, these data establish that peripheral immunosuppression observed in GBM is complex, multifaceted, and affects many immune organs. Yet, given the reproducibility of the immunosuppressive phenotype, these data establish that our model is clinically relevant in studying peripheral immunosuppression in GBM. This further allows us to investigate upstream mediators and mechanisms of immunosuppression in GBM, and to determine the relationship between distinct facets of immunosuppression reported here.

### A circulating factor(s) in glioma-bearing mice is sufficient to induce several, but not all, hallmark features of immunosuppression in naïve mice

To determine the potential role of a blood-derived factor in inducing immune suppression, we employed parabiosis. In this experiment, circulation between a naïve and glioma-bearing mouse was assessed for the capacity to induce hallmark features of immunosuppression in the conjoined tumor-free parabiont. We first surgically joined a GFP-expressing C57BL/6 mouse to a wild type C57BL/6 animal. Sharing of blood circulation was confirmed both by gross visualization of shared blood vessels as well as presence of GFP^+^ cells in the blood of both parabionts prior to tumor implantation and throughout the duration of the experiment (Figure 6 A and Figure S3 F, and Figure S4 C). One month post parabiosis surgery, the GFP-expressing mouse was intracranially inoculated with GL261 glioma cells (Figure 6A). The non-metastatic nature of this tumor was confirmed by bioluminescence imaging (Figure 6 B and Figure S3 A). Interestingly, having two conjoined immune systems did not induce early tumor rejection, as parabiotic mice became moribund at the same rate as individual mice harboring GL261 gliomas (Figure S3 B). We next asked if hallmark features of GBM (low CD4 T cell counts and MHC class II downregulation, thymic involution, and sequestration of T cells in the bone marrow) were present in parabiotic mice similar to WT mice. Our analysis revealed that peripheral CD4 T cell counts and MHCII expression levels on blood CD45^+^ cells were reduced in both naïve and glioma-bearing parabionts as a function of tumor burden (Figure 6 C-D and Figure S3 C-E).

**Figure 6:**
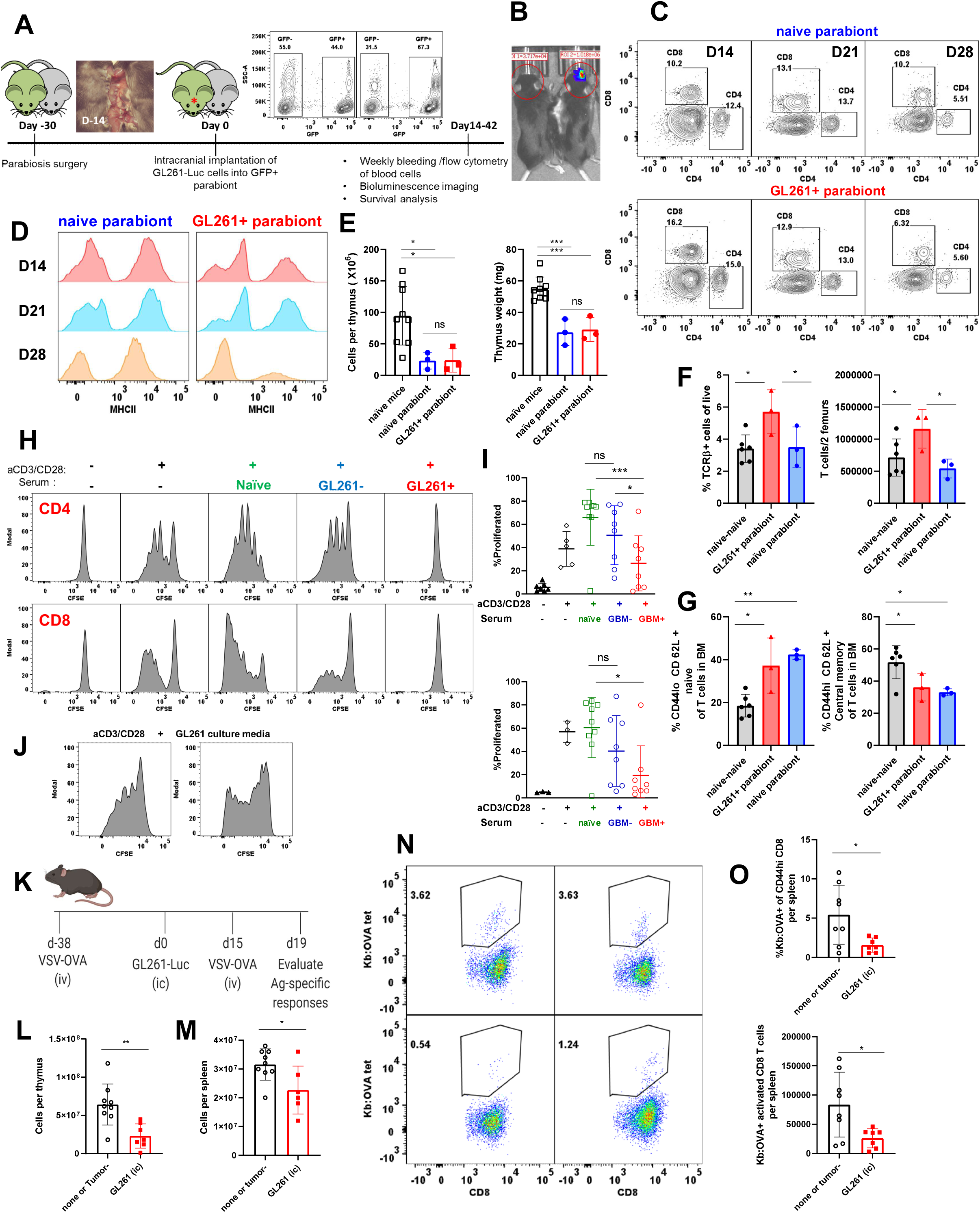
Multifaceted systemic immunosuppression in glioma-bearing mice is linked through immunosuppressive soluble factors in circulation. Schematic of parabiosis and verification of sharing of vasculature and GFP^+^ cells is shown (A). Tumor burden is shown in the tumor-bearing parabiont compared to the tumor-free partner (B). A representative flow plot of blood CD4 and CD8 frequencies is shown in both parabionts from three consecutive weeks post tumor implant (C). MHCII expression on CD45^+^ cells in the blood is shown from three consecutive bleeds (D). Thymi from naïve and glioma bearing parabionts are compared to a range of age-matched naïve controls (E). Thymic cellularity (E left) and thymic weight (F right) in glioma bearing and tumor free parabionts is compared to naïve age-matched controls. Parabiosis of a tumor-bearing to a non-tumor-bearing mouse revealed that T cell sequestration in the bone marrow only occurs in the GL261+ parabiont and not in the attached tumor free parabiont (F). However, phenotype of bone marrow resident T cells changes in both tumor-bearing and tumor-free parabionts when compared to naïve-naïve counterparts (G). Exposure to the circulation of tumor-bearing mouse is sufficient to induce enrichment in naïve L selectin+ bone marrow resident T cells in the attached animal with no glioma (G). Representative histograms depict CD4 (top) and CD8 (bottom) T cell proliferation with and without anti-CD3/CD28 Dynabeads in the presence and absence of serum from naïve, tumor ^-^ and GL261 glioma-bearing mice (G). CFSE dilution is shown (H). Percent CFSE dilution is quantified as a measure of proliferation for each condition (I). (I top) shows quantification for CD4 T cells and (I bottom) indicates CD8 quantification. GL261 cell culture media is shown to not prevent T cell proliferation in (J). Experimental design is shown for evaluation of memory responses generated against VSV-OVA during glioma progression (K). Thymic (L) and splenic (M) cellularity decreased in glioma-bearing mice previously infected with VSV-OVA compared to mice with no brain injury or tumor ^-^ controls. Frequencies and numbers of K^b^:OVA tetramer + CD8 T cells are quantified in spleens of glioma-bearing and non-glioma implanted mice 4 days post VSV-OVA re-challenge (N-O). One way Anova with Tukey’s multiple comparisons test was used to assess statistical significance for groups larger than two. For comparisons between two groups, Mann Whitney U test was used. Ns p ≥ 0.05, * p= 0.01 to 0.05, ** p=0.001 to 0.01, ***p=0.0001 to 0.001, **** p< 0.0001.

Next, we asked if thymic involution occurred in both thymi as a result of sharing circulation. Thymi recovered from both glioma bearing and tumor-free parabionts were significantly smaller as evidenced by both lower weight and cellularity compared to age-matched naïve and naïve-naïve parabiont controls (Figure 6 E and Figure S4 A-B). Parabiosis of naïve to naïve mice did not results in thymic involution, decline of blood CD4 and CD8 T cells, or MHCII expression on CD45^+^ cells in blood (Figure S4 D-E). Together, these data demonstrate that exposure to the circulation of a GL261 tumor-bearing mouse is sufficient to induce thymic involution as well as several other facets of immunosuppression (reduction in T cell count and MHCII expression) in a naïve mouse in the absence of brain injury.

### Sequestration of T cells in the bone marrow of glioma-bearing mice is not mediated through soluble factors

We sought to determine if the bone marrow specific facet of immunosuppression during neurological injuries (increased T cell sequestration as reported in Figure 5 A-B) is transferable via soluble factors. Recently, T cell sequestration within the bone marrow was put forward as a mechanism of severe immunosuppression during CNS cancers (*2*). However, the modest increase in numbers of sequestered T cells within the bone marrow does not account for the large numbers of T cells lost from the thymus, spleen and peripheral blood. T cell infiltration into the bone marrow also does not explain the loss of MHCII expression on peripheral blood cells following neurological injury. We therefore performed parabiosis and evaluated frequencies and total numbers of T cells in the bone marrow of glioma-bearing verses tumor-free parabionts.

These animals were compared to healthy conjoined parabionts. As shown in Figure 6F, we determined that T cell sequestration in the bone marrow only occurs in the glioma-bearing parabiont and not in the attached naïve parabiont. We next evaluated the phenotype of bone marrow resident T cells in both sets of parabionts. Interestingly, the bone marrow of mice with no brain tumor are comprised mainly of CD44^hi^ CD62L^+^ central memory T cells with naïve T cells as a minor population (Figure 6 G). Remarkably, we found a reduction in central memory T cells concurrently with increased frequencies of naïve T cells in bone marrows of both glioma-bearing and tumor-free parabionts when compared to naïve-naïve parabiotic pairs (Figure 6 G). This shift towards retention of naïve T cells in the bone marrow was previously shown in Figure 5 as well. Hence, exposure to the circulation of tumor-bearing mice is sufficient to induce a shift towards naïve T cells and a reduction in central memory T cells. This occurred despite the absence of increased T cell sequestration in the bone marrow. Together, these data indicate that T cell sequestration in the bone marrow as a facet of immunosuppression in the GL261 model is independent of a circulating factor(s).

### Serum isolated from mice harboring GL261 gliomas suppresses T cell proliferation *in vitro*

Parabiosis revealed that soluble factors in the circulation are sufficient to induce several features of peripheral immune suppression observed during brain injury. Next, we asked if serum obtained from GL261 glioma-bearing mice suppresses T cell function *in vitro*. We cultured CFSE labeled T cells isolated from spleens and lymph nodes of naïve mice in the presence of anti-CD3/CD28 beads which induce strong antigen independent proliferation. This stimulation results in robust activation and proliferation of both CD4 and CD8 T cells which is evident by dilution of the CFSE dye (Figure 6 H-I). To these cultures we added 5% serum isolated from naïve, GL261 tumor-bearing, or tumor ^-^ mice. T cell activation and proliferation was measured 72 hours post stimulation. Addition of serum from GL261 glioma-bearing mice (but not from naïve or tumor ^-^ mice) inhibited T proliferation *in vitro* (Figure 6 H-I). Finally, to distinguish between tumor-derived or host-derived factors, we tested whether the GL261 culture media is capable of inhibiting T cell proliferation in a similar assay. Supernatant obtained from GL261 culture had no effect on T cell proliferation suggesting that this suppressive factor is likely host-derived and not tumor-derived (Figure 6 J).

### Glioma-bearing mice are functionally immunosuppressed *in vivo*

Given that serum isolated from glioma-bearing mice potently inhibited T cell function in vitro, we next sought to determine if GBM-bearing mice had a measurable defect in T cell function *in vivo*. We focused on responses of pre-existing memory T cells in a glioma-bearing mice exhibiting multifaceted immunosuppression compared to controls. We asked whether pre-existing memory responses that were generated in the absence of brain injury were fully functional during an on-going neurological insult. To assess memory responses, we infected mice with a VSV vector bearing OVA antigen (VSV-OVA) intravenously 6 weeks before any further manipulation. We then implanted GL261 tumors in groups of mice and allowed tumors to grow (Figure 6 K). On day 15 post tumor implantation, we re-challenged with VSV-OVA and euthanized mice 4 days post re-challenge (Figure 6 K). We evaluated antigen specific responses of CD8 T cells (as a measurement of memory T cell reactivation at this time point) using tetramers against OVA (K^b^:OVA). We found that glioma-bearing mice with previous memory to VSV-OVA exhibited a reduction in thymic and spleen cellularity which was similar to non-VSV-OVA-immune glioma-bearing mice (Figure 6 L-M). Quantification of K^b^:OVA tetramer positive CD8 T cells revealed a significant reduction in frequencies and numbers of OVA specific CD8 T cells responding to reinfection in spleens of glioma-bearing mice compared to those without brain injury (Figure 6 M-O). Separately, we evaluated naïve T cell responses in glioma-bearing and tumor ^-^ mice generated against Theiler’s Murine Encephalomyelitis Virus bearing OVA antigen (*43*) (TMEV-OVA). At 7 days post intraperitoneal TMEV-OVA infection, we observed reduced CD8 T cell responses (as measured by quantifying frequencies and numbers of tetramer positive cells against both OVA (K^b^:OVA) and the immunodominant TMEV peptide, VP2_121-130_, presented in the context of the H-2D^b^ class I molecule (*73*)) in the spleens of glioma-bearing infected mice compared to tumor ^-^controls (data not shown). Together, these data provide evidence of glioma-induced immunosuppression *in vivo* that affects functions of T cells including naïve and memory antiviral responses.

### Thymic involution can be induced in multiple neurological disease models and is not unique to CNS cancer

Experimental models of CNS cancer induce sustained peripheral immunosuppression which affects several immune organs, aspects of which can be mediated through a soluble factor(s). We sought to determine the effect of acute neurological insults on the thymus. To test the effect of acute neurological injuries on thymic homeostasis, we used several models of brain insults. These models included viral infection of the brain, sterile inflammatory injury, physical injury, and seizure inductions (Figure 7 A). The brain is the primary site of insult in all these models. Therefore, effects observed in thymic homeostasis will have originated from the brain insult. To induce a viral infection in the brain, we intracranially (i.c) infected C57BL/6 mice or SJL mice with 2 x10^6^ PFU of the neurotropic virus Theiler’s murine encephalomyelitis virus (TMEV). TMEV can be cleared effectively in C57BL/6 mice whereas SJL mice experience chronic infection accompanied with demyelination (*16*). TMEV infection resulted in significant and severe thymic involution measurable by a reduction in thymic weight and cellularity seven days post infection in both C57BL/6 and SJL mice (Figure 7 B). We next investigated if generalized inflammation in the brain was capable of inducing thymic involution by intracranially injecting 1 µl of 10 µg/µl Lipopolysaccharide (LPS) into the brain and analyzing the thymus seven days later. We found significant thymic involution in LPS injected mice compared to naïve unmanipulated controls (Figure 7 C). We next tested the impact of a mild physical brain injury on the thymus. To induce physical injury, we intracranially injected 2µl of sterile PBS into the brain of C57BL/6 mice. We observed significant thymic involution by gross analysis, and reduced thymic weight and cellularity in both male and female mice seven days post injection (Figure 7 D-F). However, surgical trauma as a result of intracranial injections may affect the thymus independently of the neurologic insult. To test whether brain insults without surgical trauma also cause thymic involution, we employed a seizure induction model following intraperitoneal injection of kainic Acid (KA). KA crosses the blood brain barrier and induces neuronal excitation in the brain that results in acute seizures (*46*). In this model, we found significant thymic involution seven days post seizure induction in both male and female mice that experienced acute seizure activity (Figure 7 G, Figure S5 A-D).

**Figure 7:**
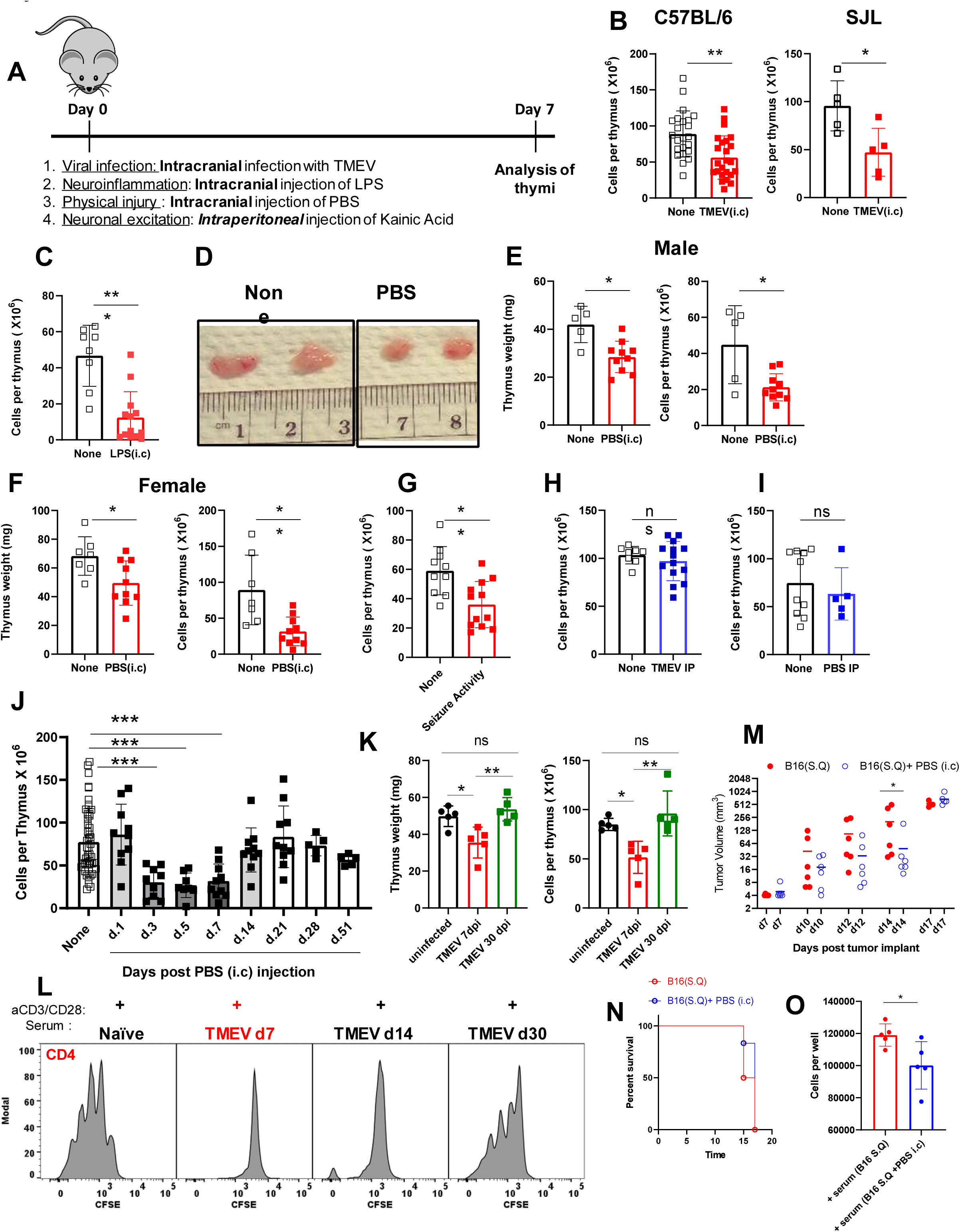
Acute neurologic insults with various origins that are delivered to the CNS result in similar thymic involution. Experimental design is shown in (A). Thymi are analyzed on day 7 post neurological insults. Thymic cellularity is quantified in C57BL/6 (B left) and in SJL (B right) mice 7days post TMEV (i.c) infection. Thymic cellularity is significantly decreased post LPS (i.c) injection compared to naïve controls (C). Gross comparison of thymi from sham control (PBS i.c) and naïve unmanipulated mice (D). Thymic weight (E left, and F left) and cellularity (E right and F right) are both significantly reduced in mice injected with PBS (i.c). The decrease in thymic cellularity post PBS injection is reproduced in both male (E) and female (F) mice. Mice injected with kainic acid that had an acute seizure activity measured by Racine’s modified scoring system, had a similar thymic involution (G). TMEV infection (H) or PBS injection (I) intraperitoneally (IP) do not induce thymic involution. Thymic involution due to intracranial injection of PBS is reversible (J). Thymi were harvested at time points post intracranial injection and cellularity was evaluated (J). Thymic involution following the clearance of TMEV is also reversible (K). Thymic involution as measured by a reduction in thymic cellularity (K left) and thymic weight (K right) is observed on day 7 post intracranial TMEV infection, but not on day 30. T cell proliferation is shown in the presence of serum isolated from TMEV (i.c) infected mice on days 7, 14 and 30 post infections (L).Growth of B16 melanoma cells in the flank of animals with brain injury (PBS injection in the brain) is slower when compared to controls (M). Survival of mice, as measured by when tumors reach end point, is similar between groups with and without brain injuy (N). B16 melanoma cells cultured in the presence of serum obtained from mice with intracranial PBS injection (d11) proliferate slower compared to serum obtained from mice without brain injury (N). Serum was collected from mice implanted with B16 melanoma (flank) on day 11 post melanoma implantation or from B16 (flank) implanted mice who also received an intracranial injection of PBS (day 4 post B16 injection) at the same time point (day 11 post melanoma injection). Melanoma tumor growth was measured using caliber. N=5-23 based on the experiment. Graphs show pooled data from 2-5 independent experiments. Data are shown as individual mice with mean. Error bars represent standard deviation. Mann-Whitney test or a one-way anova with Tukey’s multiple comparison test was used to assess statistical significance. Ns p ≥ 0.05, * p= 0.01 to 0.05, ** p=0.001 to 0.01, ***p=0.0001 to 0.001, **** p< 0.0001.

We next tested whether viral infections and physical insults in the periphery induce thymic involution. Intraperitoneal TMEV infection or PBS injection did not induce thymic involution (Figure 7 H-I), demonstrating a requirement for direct injury to the brain to induce thymic involution in these models. Overall, these data demonstrate that various acute neurological insults cause thymic involution and this form of immunosuppression is not an exclusive feature of CNS cancers alone.

### Thymic involution following acute neurological injuries is reversible upon clearance of the brain insult

To determine whether thymic involution due to acute brain injury was reversible, we used two acute injury models. First, we quantified thymic involution over time following physical injury to the brain via intracranial PBS injection. Second, we evaluated the thymic response following intracranial TMEV infection in C57BL/6 mice before and after viral clearance. Because we did not expect the intracranial PBS injection to have long lasting effects in the brain, we hence anticipated that the effect of injections on the thymus would only be observed transiently. We quantified thymic cellularity at various time points post PBS injection in the brain. Following intracranial PBS injection, we determined that thymic involution was present acutely, but was completely reversed by three weeks post injury (Figure 7 J). Additionally, as mentioned before, RNA-Seq analysis revealed that gene expression patterns of thymi isolated from mice 3-5 weeks post intracranial PBS injection did not significantly differ from naïve controls confirming reversibility of thymic immunosuppression (Figure 3). Next, we assessed thymi during TMEV infection in C57BL/6 mice. C57BL/6 mice clear TMEV infection within 21-45 days post infection (*15, 16, 20, 48*). Because C57BL/6 mice mount an effective antiviral CD8 T cell response, this model allows us to determine whether thymic involution is reversible upon viral clearance (*20, 74*). Thymic involution was not detectible in mice 30 days post TMEV infection and was only observed acutely (Figure 7 K). Together, these data suggest that thymic involution due to acute neurological injury is reversible upon clearance of the brain insult.

### Sera of mice with on-going acute neurological insults inhibit T cell proliferation

We determined that sera from glioma-bearing mice suppress T cell proliferation *in vitro*. Therefore, we next assessed whether sera isolated from mice with other neurological insults also suppress T cell function. We cultured naïve T cells with anti CD3/CD28 beads with and without serum isolated from mice post TMEV infection. Mice injected with PBS (i.c), or B16 melanoma (i.c) were also assessed using this assay. Serum obtained from TMEV infected mice at acute time points (7 dpi), but not at time points post viral clearance (30 dpi), potently inhibited T cell activation and proliferation *in vitro* (Figure 7 L, and S6 B-C). Similarly, serum acquired from PBS injected mice at acute time points was immunosuppressive while serum at times where thymic recovery was confirmed did not inhibit T cell proliferation (Figure S6 D). Serum obtained from mice implanted with B16 melanoma had immunosuppressive properties as well (Figure S6 A). These data suggest that at the time of confirmed thymic involution, serum harbors a potent immunosuppressive factor capable of inhibiting T cell activation and proliferation *in vitro*. Importantly, this factor is no longer present in serum following resolution of the neurologic insult.

### Serum-derived immunosuppression in mice with an on-going brain insult inhibits growth and proliferation of non-immune cells *in vivo and in vitro*

We next sought to further determine the extent and consequences of peripheral immunosuppression in mice with acute neurological insults. We aimed to test the growth and proliferation of non-immune cells in the presence of brain injury and soluble immunosuppressive circulating factors. We implanted mice with B16 melanoma in the flanks and allowed 4 days for tumors to grow. We then intracranially injected PBS into groups of mice and measured the melanoma growth at the peak of immunosuppression (7-14 days post injection). We found a slower growth curve for melanoma tumors in mice intracranially injected with PBS compared to controls (Figure 7 M). The slower growth of melanoma tumors occurred at the peak of immunosuppression in this model as growth reached the levels observed in controls by day 17 (Figure 7 M). This slower growth of melanoma tumors was likely not due to better anti-tumor immune responses for two reasons. First, we did not observe an overall survival difference in these mice or differences in immune responses measured at a matched time point (Figure 7 N and data not shown). Second, we tested the direct growth of melanoma cells *in vitro* in the presence of sera isolated either from mice with B16 (flank) tumors or sera isolated from mice with B16 (flank) tumors that also received an intracranial PBS injection (Figure 7 O). We observed a significant reduction in proliferation of melanoma cells exposed to sera from mice with a recent brain injury compared to those without *in vitro* (Figure 7 O). Together, these data suggest that serum of mice with on-going brain injury harbors a potent factor that can inhibit proliferation of both immune and non-immune cells and affect outcomes of immunological challenges outside of the CNS.

### Adrenal glands play a crucial role in maintaining immune organ size at baseline and during neurological insults

We next aimed to elucidate whether cortisol and other stress hormones play a significant role as soluble mediators of peripheral immunosuppression that affect the thymus, spleen, peripheral blood, and the bone marrow. While the thymus analysis both by RNA-seq and via flow cytometry did not strongly support cortisol as the sole mediator of thymic involution, we sought to determine the level of peripheral immunosuppression affecting immune organs during sustained neurological insults in adrenalectomized mice. Cortisol is produced by adrenal glands under the influence of the pituitary (*57*). We purchased commercially-available adult adrenalectomized mice from Jackson Laboratory. Adrenalectomy was verified by Jackson Laboratory before shipment of mice. We sought to determine the level of immunosuppression affecting blood, spleen, and thymus in the absence of adrenal glands following GL261 implantation in the brain. Interestingly, we found that adrenalectomy alone is sufficient to increase blood, thymus, and spleen cellularity (Figure 8 A-C) in naïve mice. This increase in total cellularity in blood, spleen, and thymus was not due to expansion of particular immune cells as frequencies of T and B cell populations appeared comparable between naïve WT and adrenalectomized mice (Figure S7 A-C and data not shown). Next, we assessed the population of bone marrow resident T cells in naïve wild type and adrenalectomized mice. Surprisingly, we found significantly fewer bone marrow resident T cells in naïve adrenalectomized mice compared to controls (Figure 8 D). These data suggest that bone marrow sequestration of T cells during homeostatic conditions is at least partially regulated by systemic cortisol levels in naïve animals. These data alone suggest that hormones produced by the adrenal glands in naïve mice play a crucial role in regulating the cellularity of immune organs. This unique homeostatic role of stress hormones has been previously underappreciated. Because of this homeostatic role of cortisol in regulating immune organ size, it may be difficult to dissect changes induced due to brain injury from those stemming from lack of the homeostatic role of these hormones. However, with that caveat in mind, we performed tumor implantations and evaluated various facets of immunosuppression in adrenalectomized mice bearing GL261 gliomas compared to controls. Analysis of the peripheral blood revealed that in contrast to GBM-bearing WT mice, adrenalectomized mice did not exhibit reductions in CD4 or CD8 T cells, or reduced MHCII expression levels on hematopoietic cells (Figure 8 E-G and S7 G-H). Similarly, analysis of thymi from glioma-bearing adrenalectomized mice revealed no evidence of thymic involution, loss of DN3 cells, DP cells, or accumulation of single positive cells compared to naïve adrenalectomized controls (Figure 8 H and S7 K-M). In spleen, we did not observe evidence of organ atrophy or loss of T cells or MHCII down-regulation in glioma-bearing adrenalectomized mice compared to controls (Figure 8 I and S7 I-J).

**Figure 8:**
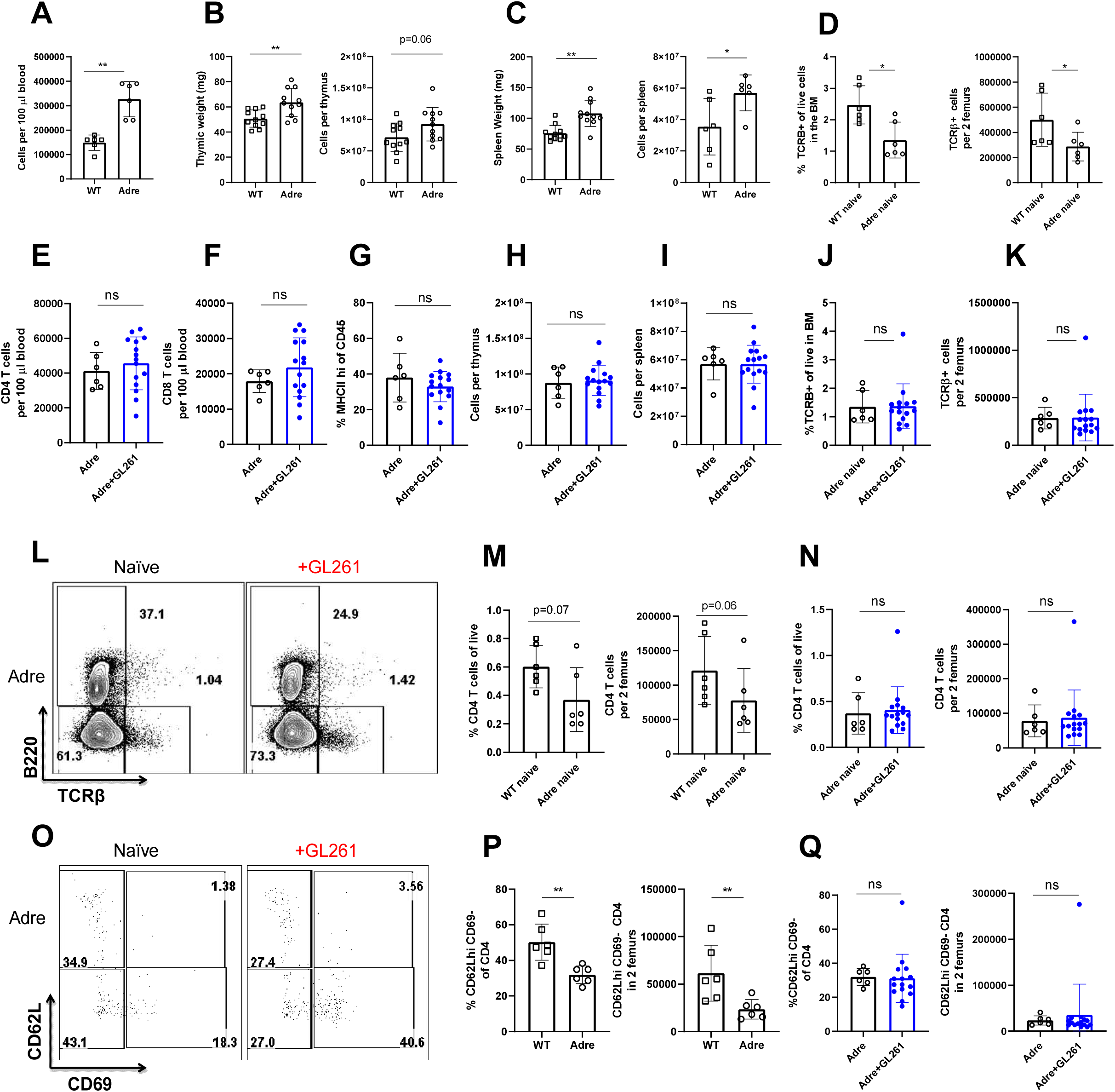
Hormones produced by the adrenal gland control immune organ size and cellularity at baseline Cellularity of blood per equal volume is increases in naïve adrenalectomized mice compared to naïve WT. (A). Thymic weight and cellularity is increased in naïve adrenalectomized mice compared to naïve WT (B). Spleen weight and cellularity is increased in naïve adrenalectomized mice compared to naïve WT (C). Bone marrows of naïve WT mice harbors an increased frequency and numbers of T cells when compared to adrenalectomized controls (D). CD4 (E), CD8 (F) and frequency of MHCII expression on CD45+ cells (G) in blood is equivalent between naïve and glioma-bearing adrenalectomized mice. Similarly, cellularity of the thymus (H) and spleen (I) does not change in glioma-bearing adrenalectomized mice compared to controls. Frequency (J) and numbers (K) of bone marrow resident T cells increase in glioma-bearing adrenalectomized mice compared to naïve adrenalectomized controls. Representative flow plot of T cells within the bone marrow of naïve and glioma-bearing adrenalectomized mice is shown (L). Fewer bone marrow resident CD4 T cells are found in naïve adrenalectomized mice compared to naïve WT mice (M).Frequencies and numbers of bone marrow resident CD4 T cells does not increase in glioma-bearing adrenalectomized mice when compared to adrenalectomized naïve controls (N). Phenotype of bone marrow resident CD4 T cells in naïve and glioma bearing mice reveals enrichment in CD62L+ cells (O). This population is significantly decreased at baseline in naïve adrenalectomized mice when compared to WT mice (P). In contrast to WT mice, no increase in CD62L+ CD69-CD4+ T cells occurs in the bone marrow of glioma-bearing adrenalectomized mice (Q). N=6-14. Individual data are shown. Error bars represent standard deviation. Mann Whitney U test was used. Ns p ≥ 0.05, * p= 0.01 to 0.05, ** p=0.001 to 0.01, ***p=0.0001 to 0.001, **** p< 0.0001.

We next evaluated changes within the bone marrow T cell compartment following GL261 implantation. The increase in bone marrow resident T cells following glioma implantation did not occur in adrenalectomized mice (Figure 8 J-L and S8 A-C) when compared to controls. It is worth mentioning here that all hallmark features of neurological insult induced immunosuppression still occurred in glioma-bearing WT mice compared to naïve WT controls as seen before (Figure 5, S7 G-M and data not shown).

Similar to trends observed in the overall T cell counts within the bone marrow, we observed fewer bone marrow resident CD4 T cells in naïve adrenalectomized mice compared to WT controls (Figure 8 M). In contrast to WT mice, total frequencies and absolute counts of CD4 T cells did not further increase in adrenalectomized glioma-bearing mice compared to naïve controls (Figure 8 N). Consistent with these trends, we also determined that naïve adrenalectomized mice harbor fewer CD62L^+^ CD69^+^ CD4 T cells in the bone marrow compared to naïve WT mice (Figure 8 O and P). While the increase in CD62L^+^ CD69^-^ CD4 T cells in the bone marrow accounted for the observed T cell accumulation in glioma-bearing WT mice (Figure 5 E-F), this population did not expand in glioma-bearing adrenalectomized mice (Figure 8 Q).

Together these data support a role for cortisol and stress hormones in maintaining immune organ size and peripheral blood cellularity at baseline. In the absence of adrenal glands, thymic and spleen involution, loss of CD4 and CD8 T cells, MHCII downregulation, and T cell sequestration in the bone marrow did not occur during GBM progression (Figure 8, S7 and data not shown).

### Serum from glioma-bearing mice harbors a high molecular weight non-steroid immunosuppressive factor

We have thus far determined that serum isolated from glioma-bearing mice is potently immunosuppressive (Figure 6 H-I). Yet, due to the homeostatic role of cortisol at baseline, we were unable to rule out the possibility of steroids as mediators of immunosuppression in serum. Hence, we sought to determine the molecular weight of the immunosuppressive molecules found in this serum. Through a series of molecular weight cut-off filtration strategies we determined that large factors with molecular weights greater than 100 kDa are immunosuppressive and capable of inhibiting T cell proliferation *in vitro* (Figure 9 A). Small molecules below 3 kDa were not immunosuppressive (Figure 9 A). This argues against steroids and small molecules as mediators of immunosuppression. We next sought to further validate that cortisol is not responsible for the immunosuppressive properties of serum isolated from glioma-bearing mice. We isolated serum from WT and adrenalectomized mice that were intracranially implanted with GL261 gliomas. Interestingly, sera isolated from glioma-bearing WT and adrenalectomized mice were equally immunosuppressive *in vitro* (Figure 9 B-C). This is while sera isolated from naïve WT or adrenalectomized mice did not affect T cell proliferation *in vitro* (Figure 9 B-C). Similarly, adrenalectomized mice did not have a survival benefit and succumb to death with similar kinetics to WT mice (these mice actually had a slightly lower survival (MST 23) compared to WT mice (MST 27) (Figure 9 D).

**Figure 9:**
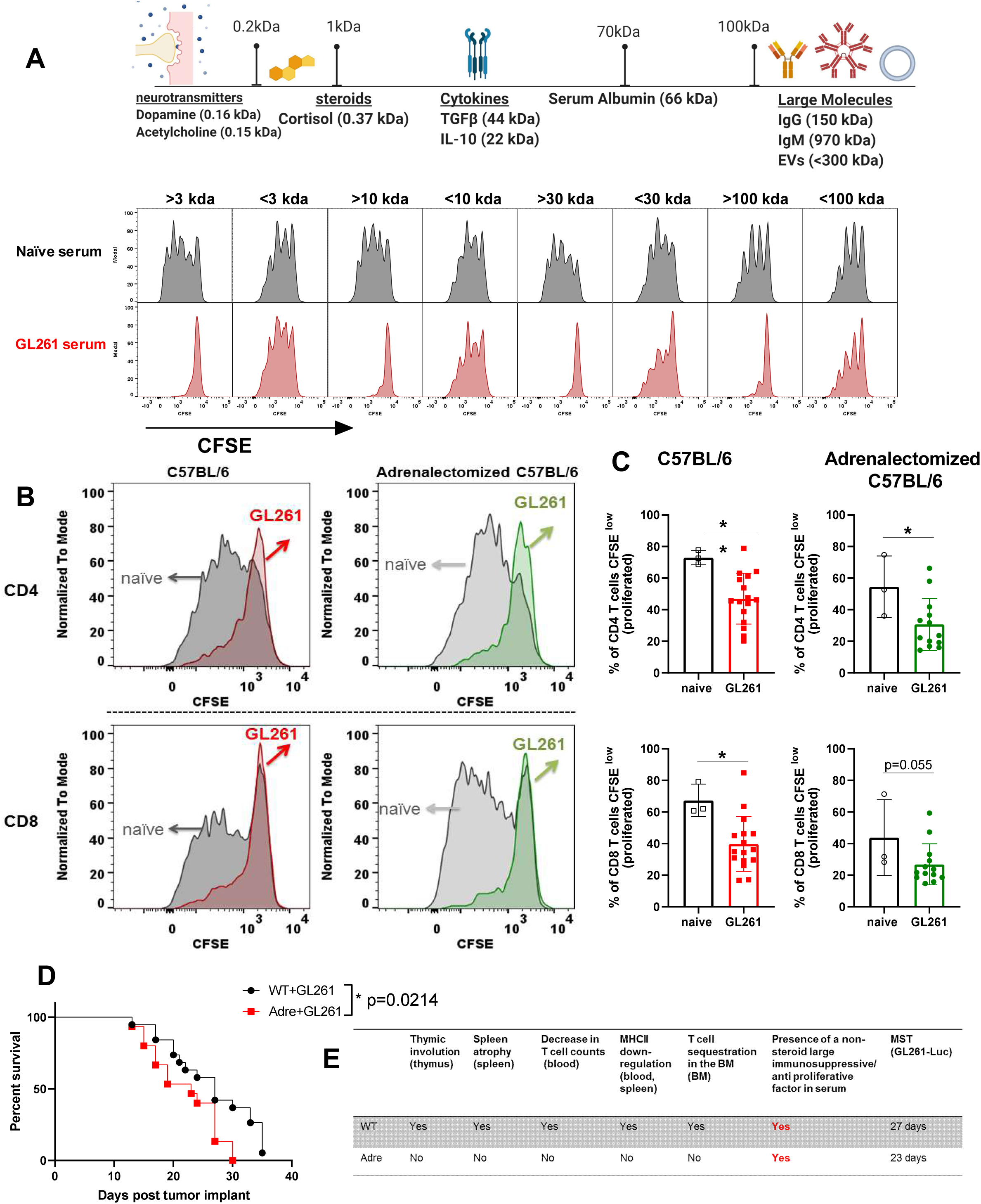
Serum of glioma bearing mice harbors a novel immunosuppressive factor with molecular weight larger than 100 kiloDaltons Serum obtained from naïve or glioma-bearing mice were isolated and passed through protein columns with a molecular weight cutoff of 3, 10, 30, or 100 kda. Both top and bottom fractions were collected and their ability to inhibit T cell proliferation was tested. T cells obtained from naïve mice were labeled with CFSE and cultured with aCD3/CD28 dynabeads in the presence of individual fractions isolated from naïve or GBM-bearing sera. Proliferation was measured 72 hours later using flow cytometry. Serum fractions with high molecular weights (>3, >10, >30, >100) were deemed immunosuppressive as they potently inhibited T cells proliferation *in vitro* (A). This data argues against cortisol or other small molecules related to stress hormone pathways playing a role as the immunosuppressive factor produced by the brain during neurological injuries. The molecular weight of the immunosuppressive factor released into serum during neurological injuries is greater than 100 kda. Consistent with this, we tested the serum obtained from glioma bearing WT or adrenalectomized mice. Serum of glioma bearing mice from both WT and adrenalectomized mice inhibited T cell proliferation *in vitro* (B-C). Glioma-bearing adrenalectomized mice do not have a survival benefit over WT mice despite lacking several facets of immunosuppression (D). Adrenalectomized mice only had the immunosuppressive factor aspect of immunosuppression and did not have other facets of immunosuppression (including thymic and spleen involution, T cells reduction in blood, and accumulation in the bone marrow, or MHCII downregulation) (E). However, this data suggest that the immunosuppressive serum-derived factor alone is sufficient to block anti-tumor responses and prevent a survival benefit. For A, 10 GL261 sera were pooled for analysis. For B-C N=3-10. Data are shown as individual mice with mean. Error bars represent standard deviation. Mann-Whitney test was used to assess statistical significance. Ns p ≥ 0.05, * p= 0.01 to 0.05, ** p=0.001 to 0.01, ***p=0.0001 to 0.001, **** p< 0.0001.

Together, our analysis found several facets and mechanisms of immunosuppression at play during neurological insults that affect immune organs. For example, we reported that adrenalectomized mice did not have thymic or spleen involution, a reduction in T cell counts or a reduced MHCII expression (Figure 8). However, serum isolated from these animals was as immunosuppressive as WT animals and these mice succumb to death with similar kinetics (Figure 9 D). This suggest that the non-steroid immunosuppressive soluble factor found in circulation of both WT and adrenalectomized mice during neurological insults is perhaps sufficient to induce a lower state of immunity, block proliferation of T cells, and inhibit anti-tumor responses (Figure 9 E). This also suggests that in the presence of this immunosuppressive factor, reversal of any of the other facets of immunosuppression observed by us and other groups (including CD4 T cells counts, thymic and spleen involution, and MHCII expression levels) alone may not directly translate into a survival benefit. Instead, focused efforts into identification of the immunosuppressive factor in circulation may help aid in both improving survival and reversal of immunosuppression observed in several immune organs.

## Discussion

In this study, we thoroughly and mechanistically analyzed both primary and secondary immune organs during neurological insults and describe at least 6 distinct facets of immunosuppression in experimental GBM: 1. thymic involution associated with a previously unappreciated unique cellular and transcriptomics signature, 2. spleen involution, 3. loss of CD4 and CD8 T cells from circulation and secondary lymphoid organs, 4. downregulation of MHCII on hematopoietic cells, 5. release of non-steroid immunosuppressive factors with large molecular weight in serum that block proliferation of cells, and 6. Major phenotypic changes in resident T cells within the bone marrow. These findings demonstrate that CNS cancers cause multifaceted immunosuppression which concurrently affects peripheral blood, serum, bone marrow, thymus, and spleen: together, this induces a state of lowered immunity. Our data further demonstrate that immunosuppression due to neurological insults is not exclusive to brain tumors as it occurs following several acute neurological insults as well suggesting this response is likely generated by the brain and not the individual insults. This study also provides detailed analysis regarding the role of stress responses, non-steroid soluble factors, and other possible mechanisms ensuing hallmark features of immunosuppression. However, further studies are needed and currently underway in our laboratory to determine whether the multifaceted brain-injury induced immunosuppression of distinct immune organs is dependent on each other.

### Thymic involution is an immunosuppressive feature of experimental GBM and other neurological injuries

Immunosuppression following a variety of neurological insults has been broadly studied both clinically and in animal models (*2, 7, 8*). In stroke patients, over one third of mortalities are due to immune suppression alone, yet the mechanism of such immunosuppression remains unknown (*8, 11, 31, 32*). In GBM, the overt immunosuppression seen in patients is attributed to tumor microenvironment and factors produced by the tumor itself (*2, 3, 5, 6*). It is worth mentioning that the thymus is reported to shrink following traumatic brain injury (TBI) and experimental GBM in mice (*2, 8, 75*). However, thymic involution following TBI and GBM has been attributed to the specific pathophysiology of these conditions, rather than an underlying intrinsic peripheral response to CNS insult. Thymic involution leads to lower T cell production and disruptions in T cell selection, leading to a lack of adequate peripheral T cell replenishment when needed. In contrast to these studies, we report for the first time that thymic involution following brain injury is a pathology-agnostic response with a unique signature, as it is a reaction by the brain and hence does not depend on the nature of injury.

We demonstrated that both acute and chronic neurological injuries of various origins cause thymic involution (Figure 1 and 7). This thymic involution was reversible upon clearance of the insult, and its magnitude depended on the extent of brain injury (Figure 1 and 7). Identical insults induced peripherally did not affect thymic homeostasis (Figure 1 and 7). These findings demonstrate the presence of a brain-thymus communication which is a general feature of neurological injuries.

Our results strongly predict that thymic involution could affect children with diverse brain tumors (including those with benign tumors), altering T cell repertoire and rendering these patients immunosuppressed for an extended period. Thus, it is crucial to study how thymic involution affects future T cell repertoire as well as T cell responses. Intriguingly, RNA-Seq analysis of our data found slight changes in several genes related to T cell development in thymi that had recovered from brain injury compared to naïve controls suggesting even transient mild brain insults can affect long term T cell developmental programs (Figure 3 and S2).

Thymic involution is similarly detrimental in adults. The thymus can sense peripheral T cell numbers and respond by increasing T cell production if numbers are low (*36*). In the absence of a sudden insult against peripheral T cells, the thymus may remain small in adults. However, the thymus does not lose its ability to increase in size and generate new T cells in the event of peripheral immunosuppression. The crucial role of the thymus in reconstituting T cell numbers in adults following immunosuppression has been demonstrated in transplant patients, in patients with HIV, and in patients post chemotherapy (*34, 36, 37, 40, 76-78*). Our study shows that thymic involution and its reversal directly reflect the extent of brain insult and is caused by soluble factors in circulation. Understanding these immunosuppressive mechanisms offers hope for future pediatric and adults patients with neurological trauma.

#### The adult thymus is uniquely affected by a CNS tumor

In this study, we demonstrated that transcriptional programming of the thymus was directly altered by a non-metastatic brain tumor in adult mice (Figure 3). We found epithelial cell development and maturation programs to be enhanced, while T cell commitment programs were decreased (Figure 3). This correlated with a block in T cell proliferation at the DN2-DN3 stage. DN2-DN3 stage of T cell development is associated with high levels of proliferation following TCRβ chain selection (*52*). A block in DN2-DN3 is likely caused by a block in proliferation which is consistent with the most downregulated pathways analyzed by RNA-seq being involved in DNA replication and cell proliferation (Figure 3). We also demonstrated that thymic involution was caused by circulating factors using parabiosis. Confirming this result, serum obtained from mice with various neurological insults potently inhibited activation and proliferation of T cells *in vitro* at the time of established thymic involution (Figure 6 and 7). Given that thymic involution can be transferred through serum-derived soluble factors released during neurological injuries that are potent inhibitors of cell proliferation, it is strongly possible that these factors intervene at the DN2-DN3 stage of T cell development. Additionally, we found an increase in B cell frequencies in the thymus which can be explained by a decrease in *Gata3* expression. Normally, *Gata3* inhibits B cell development and lineage commitment in the thymus (*51–53, 79*). Hence, it is conceivable that during neurological injuries a decrease in *Gata3* translates into an enhanced B cell development program.

We also found an increase in total single positive cells and TCRβ^+^ single positive CD4 and CD8 T cells in the thymus (Figure 2). Interestingly, we found a significant downregulation in the expression of *Coro1a* gene (Figure 3). This gene is extremely important for T cell exit and sensitivity to S1P both in the thymus and secondary lymphoid organs (*71*). The downregulation of *Coro1a* in the thymus can explain the accumulated single positive phenotype as Coro1a knockout mice exhibit accumulated single positive cells in the thymus and have severe peripheral lymphopenia (*71*). It is also interesting to note that recently downregulation of S1PR1 during CNS cancers was deemed responsible for sequestration of T cells in the bone marrow as part of the overall immunosuppression (*2*). This perhaps warrants future studies into mechanisms of *Coro1a* downregulation in the thymus (and also in spleen) during neurological insults and how it relates to S1P sensitivity.

#### Thymic involution resulting from experimental GBM presents with a unique pattern which is distinct from stress induced involution

Thymic involution and stress responses have been linked for many decades(*55*). We found a significant decrease in DP cells in the thymus while single positive cells appeared to be accumulating within the thymus (Figure 2). This is interesting as a decrease in DP cells (mainly through enhanced cell death pathways) can be explained by high cortisol levels due to stress responses (*55, 57, 59, 61*). However, the brain-injury induced effect on DP cells is likely not due to stress hormones alone because of the following reasons. 1. We did not observe enhancement or enrichment of genes involved in stress responses or genes involved in glucocorticoid receptor activation in our RNA-seq data set. 2. We did not observe activation or enrichment of genes associated with the glucocorticoid pathway in our RNA-seq data set or through flow cytometry. 3. Concurrently with a loss in DP cells, we observed a block in DN2-DN3 stages of development as well as accumulation of single positive T cells and enhanced B cell generation in the thymus. Neither a DN2-DN3 block, nor a single positive T cell accumulation, or an enhanced B cell development signature have been reported to be associated with stress induced, infection induced, or age mediated thymic involution (*55, 57-61, 72, 80, 81*). Surprisingly however, we found that the adrenal gland plays a crucial role in maintaining cellularity and weight of thymus and spleen at baseline (Figure 8). It also plays a role in maintaining overall numbers of T cells in blood and bone marrow (Figure 8).

#### The thymus involutes with brain injury and returns to baseline homeostasis after the injury is cleared demonstrating thymic plasticity

This is seminal for two reasons. First, the thymus can detect a brain injury and accordingly respond through involution. Second, this thymic involution is reversible. The thymus successfully returns to a pre-insult state if the brain injury clears, demonstrated by both the recovery of thymic cellularity as well as an enriched recovery signature measured through transcriptional programming (PBS vs. naïve group Figure 1M, Figure 3 and Figure S2). Understanding mechanisms of thymic involution as well as reversal are both important. Inducing thymic reversal may be useful in preventing T cell suppression during CNS cancers, and also during aging where thymic regeneration in the absence of peripheral immunosuppression may be triggered to improve responses to vaccinations and pathogens in the elderly. The reversibility of the thymic involution in our work allows identification of potential targets for thymic regeneration.

### Immunosuppression affecting blood and spleen are additional facets of immunosuppression induced by neurological injuries

We found a significant loss of CD4 T cells, CD8 T cells and MHCII expression on hematopoietic cells in the peripheral blood and spleen (Figure 4). These immunosuppressive features occur concurrently with changes in the thymus and can also be transferred through soluble factors. This suggests that systemic release of large molecular weight immunosuppressive factors affects not only T cell development in the thymus, but also T cell activation, clonal expansion and maintenance in the periphery. This state of immunosuppression is extremely detrimental because a continuous loss of T cells from secondary lymphoid organs and blood concurrent with lack of replacement either through homeostatic proliferation of memory T cells or through generation of new T cells from the thymus results in a sustained state of immunosuppression. For example, we demonstrated through infection studies using VSV vectors that even previously generated memory T cells exhibit decreased responses in the presence of brain injury (Figure 6). This suggests both homeostatic proliferation and pre-existing memory responses will likely be diminished in patients with neurological diseases.

### T cell sequestration in the bone marrow has a complex mechanism

We reproduced bone marrow sequestration of T cells during neurological insults in our model of GL261 similar to a recent report in other models of CNS cancers (Figure 5 and 8 and (*2*)). While a modest increase in bone marrow resident T cells is evident during neurological injuries, this population mainly consists of CD62L^+^ CD69^-^ CD44^+^ and CD44^-^ naïve and central memory CD4 T cells (Figures 5, 6 and 8). CD62L^+^ CD69^-^ CD44^+^ and CD44^-^ naïve and central memory CD4 T cell populations are present in the bone marrow of both naïve and glioma harboring mice. These populations are significantly diminished in naïve adrenalectomized mice implying that cortisol plays a homeostatic role in maintaining the size of the T cell pool in the bone marrow as well. Our parabiosis experiments clearly demonstrated that while the sequestration of bone marrow T cells only occurs in the glioma-bearing parabiont, a change in phenotype of bone marrow resident cells towards naïve phenotype was evident in both parabionts compared to naïve-naïve parabionts (Figure 6). This suggests that soluble factors (released during glioma growth) affect the bone marrow niche and induce a change that is reflected in a shift towards naïve T cells within the bone marrow. This is while the overall T cell sequestration during neurological insults is likely due to neuronal connections between the brain and bone marrow and not due to circulating factors. Importantly, this mild sequestration of T cells within the bone marrow cannot explain all other facets of immunosuppression discussed earlier including thymic changes, severe loss of T cells from blood and spleen, anti-proliferative effects of serum on immune and non-immune cells, or lack of responsiveness of previously activated memory T cells upon re-challenge.

### The modest increase in bone marrow resident naïve CD4 T cells does not account for severe defects observed in all immune organs

It has been put forward that T cell sequestration in the bone marrow is a component of experimental GBM induced immune suppression(*2*). We too observe this in our studies. However, mathematically we contend that this sequestration does not account for the magnitude of the multifaceted immune suppression we observe. We calculated mean thymic cellularity, spleen cellularity, numbers of CD4 T cells per 100 µl of blood, and numbers of accumulated T cells in the bone marrow of naïve and GL261-bearing mice to obtain values that represent average cell loss/gain within our experiments. Comparing naïve to glioma-bearing mice, 57 million cells from the thymus, and 18 million cells from the spleen were lost. In spleen alone, CD4 T cells declined on average by 3.8 million cells. Similarly, blood CD4 T cell compartment was reduced by about 7900 cells per 100 µl volume of blood. Assuming a mouse has about 1 to 1.5 ml of blood by volume, we can estimate an approximate loss of 80000-83000 CD4 T cells from blood alone. In contrast to these significant losses within the T cell compartments of thymus, spleen, and blood, the bone marrow T cell compartment only expanded on average by 286000 cells. The gain in bone marrow resident CD4 T cells was equivalent to about 140000 cells. We realize that accounting for changes within two femurs does not fully recapitulate numbers in the entire bone marrow compartment in the mouse. Nevertheless, loss of 57 million thymocytes, 3.8 million splenic CD4 T cells, and 83000 blood CD4 T cells cannot be explained through a modest increase of 140000 CD4 T cells in the bone marrow. Additionally, all T cells, including naïve and memory CD4 and CD8 T cells, are lost to about equal levels from the spleen and blood, yet we mainly observed an enrichment in bone marrow resident naïve CD4 T cells. Finally, we did not observe evidence of premature exist of thymocytes from the thymus, nor did we observe accumulation of T cells with a DP phenotype in blood or bone marrow. Through the above analysis, we put forward that bone marrow sequestration of resident T cells is not the sole mechanism of neurologic-injury induced immunosuppression. We also propose that changes in circulating factors induced by brain injury account, at least in part, for alterations in primary and secondary immune organs.

### Systemic immunosuppression caused by CNS-cancers can be potentially restored through targeting circulation

This study creates a path forward to reverse systemic immunosuppression through targeting of serum derived factors. Interestingly, adrenalectomized mice implanted with GL261 glioma did not have thymic involution, spleen atrophy, a clear evident loss of T cells, downregulation of MHCII expression, or sequestration of T cells within the bone marrow. They, however, harbored the potent immunosuppressive factor(s) in their serum and this alone translated into similar anti-tumor responses and survival times compared to WT mice (Figure 9 and data not shown). Therefore, we hypothesize that reversing immunosuppression mediated through the serum-derived factors should be the first step in reversing immunosuppression. This hypothesis is supported by the evidence obtained from adrenalectomized mice in which reversal of all facets of immunosuppression in the presence of the potent immunosuppressive factors in serum did not achieve a survival benefit (Figure 9 D-E). In the absence of therapeutic targets aimed at reversing serum-derived immunosuppression, targeting other facets of immunosuppression alone may fail. Hence, reversal of immunosuppression in serum could be the first step in reversing systemic immunosuppression as it will likely lead to enhanced thymic development. This in turn will enhance T cell numbers in the periphery. In parallel, removing the anti-proliferative factor from the circulation will potentiate homeostatic proliferation and T cell activation while boosting naïve and memory T cells responses. Consequently, this would enable better endogenous anti-tumor responses and improve immunotherapy outcomes. Additionally, lifting the inhibition on T cell proliferation can enhance pre-existing memory responses, and induce better naïve T cell priming against novel antigens. Identifying serum-derived non-steroid immunosuppressive factor(s) is therefore crucial in understanding and reversing immunosuppression during neurological insults to improve current and future therapies for the large cohort of patients with GBM, as well as for those with other neurological diseases.

## Acknowledgements

We would like to thank Drs. Robert L Fairchild, Ian Parney, Roberto Cattaneo, and Steven Rosenfeld for their crucial help and support in preparing this manuscript. We would like to thank Drs. Barsha Dash, Courtney S Malo, and Sinéad Kinsella for their technical assistance, help in designing experiments, and conceptualizing the manuscript. We would like to also thank Mayo Clinic’s sequencing and bioinformatics cores for technical support. We also like to thank the NIH tetramer facility for providing tetramers that were used in this study. Work presented in this manuscript was partially funded through the following sources: NINDS 1R01 NS 103212-01 (AJJ), NIH T32 training grant through Mayo Clinic Neuro-Oncology program (KA), Brains together for a cure foundation (KA), Mayo Clinic internal grant funding through the Department of Molecular Medicine small grants (KA), Center for MS and Autoimmune Neurology fellowship (KA), and Immuno-oncology program (AJJ and KA).

---

**Figure S1:**
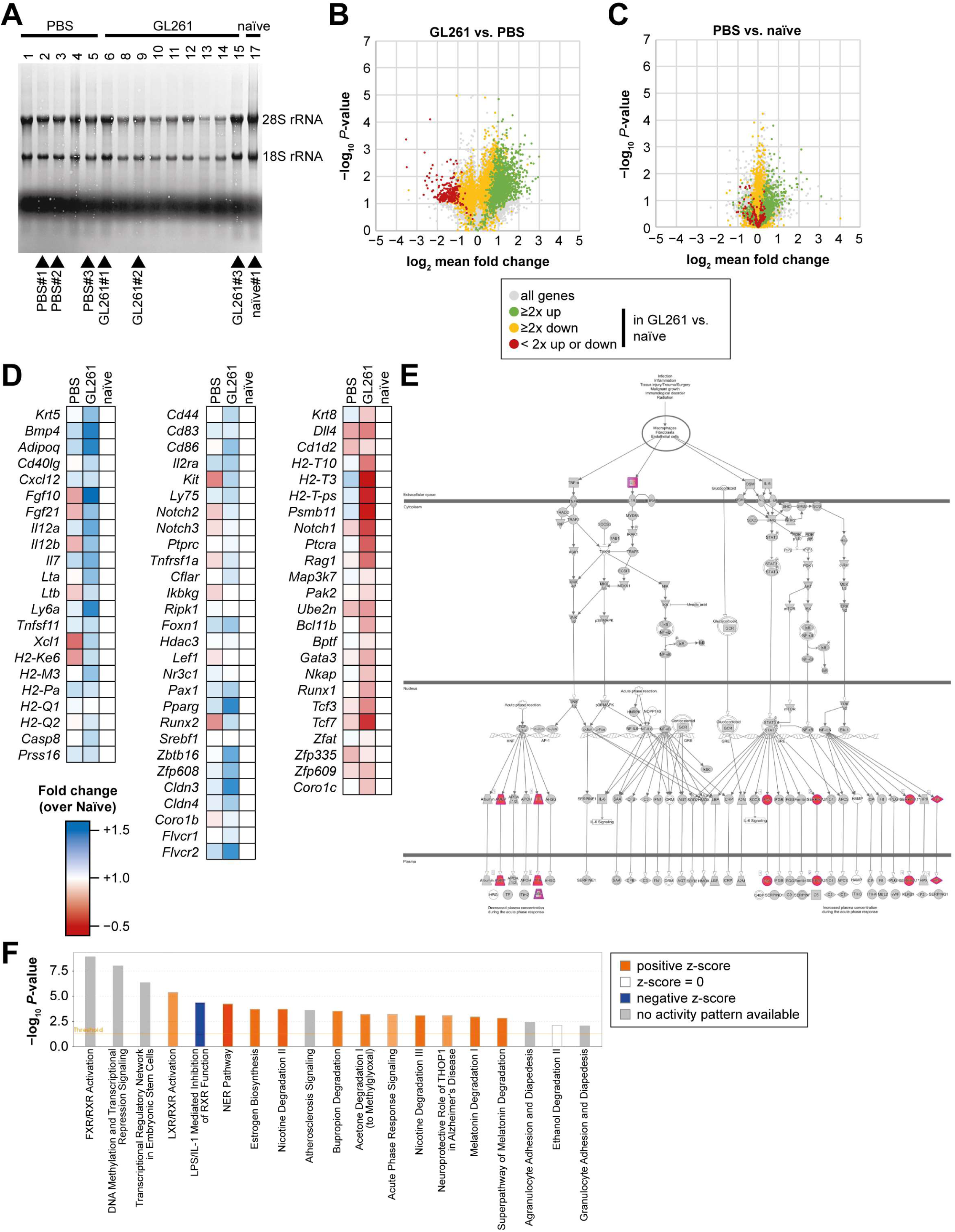
GL261 glioma-induced changes in the thymus are distinct from those induced by stress: Quality of isolated RNA from thymi was tested using methylene blue staining (A). Differential gene expression in GL261 glioma bearing mice is compared to sham and naïve controls (C-D). For statistical analysis, we used a one-way ANOVA for every gene. In green are genes that are significantly upregulated in GL261 vs Naive by at least 2 folds. In red are respectively downregulated at least 2x and significant by ANOVA. Yellow genes are regulated less than 2x up or down, but still significantly regulated as evaluated using ANOVA. Grey genes are not significantly regulated in ANOVA, even though they may or may not be more than 2x regulated. (D) Shows expression of genes that are at least 2 fold up or downregulated amongst our gene set between GL261-bearing and naïve control thymi, but did not reach statistical significance using ANOVA. Genes significantly regulated in the thymus of GL261 glioma bearing mice compared to naïve controls that belong to the acute phase response pathway (E). Acute phase response pathway is amongst top highly enriched pathways in our data set. Predictions of most significant pathways enriched in our thymic RNA-seq data set according to IPA (F).

**Figure S2:**
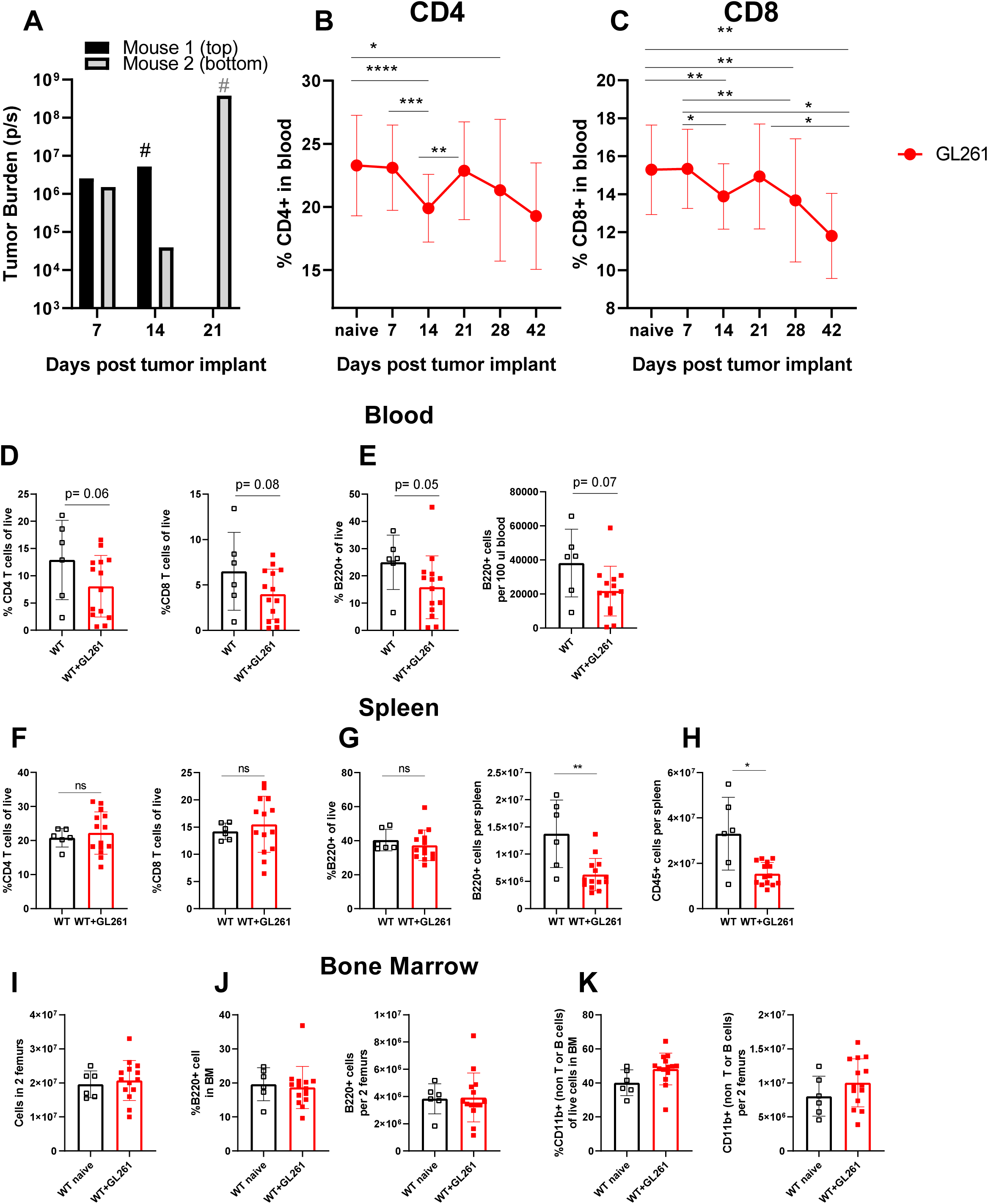
A sustained and continuous decline in T cells is a hallmark feature of a growing glioma in the brain and only transiently occurs following acute insults. Tumor burden is tracked in representative mice shown in Figure 4 A-B in (A). # Represents the day mouse succumb to death (14 and 21 for the representative mice shown). Frequencies of CD4 (B) and CD8 (C) are shown in mice implanted with GL261 cells. Frequencies of CD4, and CD8 (D) and B cells (E left) are shown as a frequency of live cells in blood. Counts of B220^+^ cells are shown per 100 ul of blood (E right). For T cell gating, we gated on singlets, live, CD45^+^ TCRB^+^ B220-cells and then gated on either CD4 or CD8 T cells. For B220 gating, we used a similar gate and instead focused non TCRβ^+^ cells that stained positive for B220. Frequencies of CD4, CD8 (F) and B220^+^ cells (G left) are shown as a fraction of live cells per spleen. Counts of B220^+^ (G right) and CD45^+^ (H) are shown. Total cellularity of bone marrow does not change between naïve and glioma-bearing mice (I). Similarly, frequencies and numbers of B cells (J) and CD11b+ cells (K) in the bone marrow does not change between naïve and glioma-bearing mice. One way Anova with Tukey’s multiple comparisons (when comparing more than two groups) test or a Mann Whitney U test was used to calculate statistical significance (when comparing two groups) was used to assess statistical significance. Ns p ≥ 0.05, * p= 0.01 to 0.05, ** p=0.001 to 0.01, ***p=0.0001 to 0.001, **** p< 0.0001.

**Figure S3:**
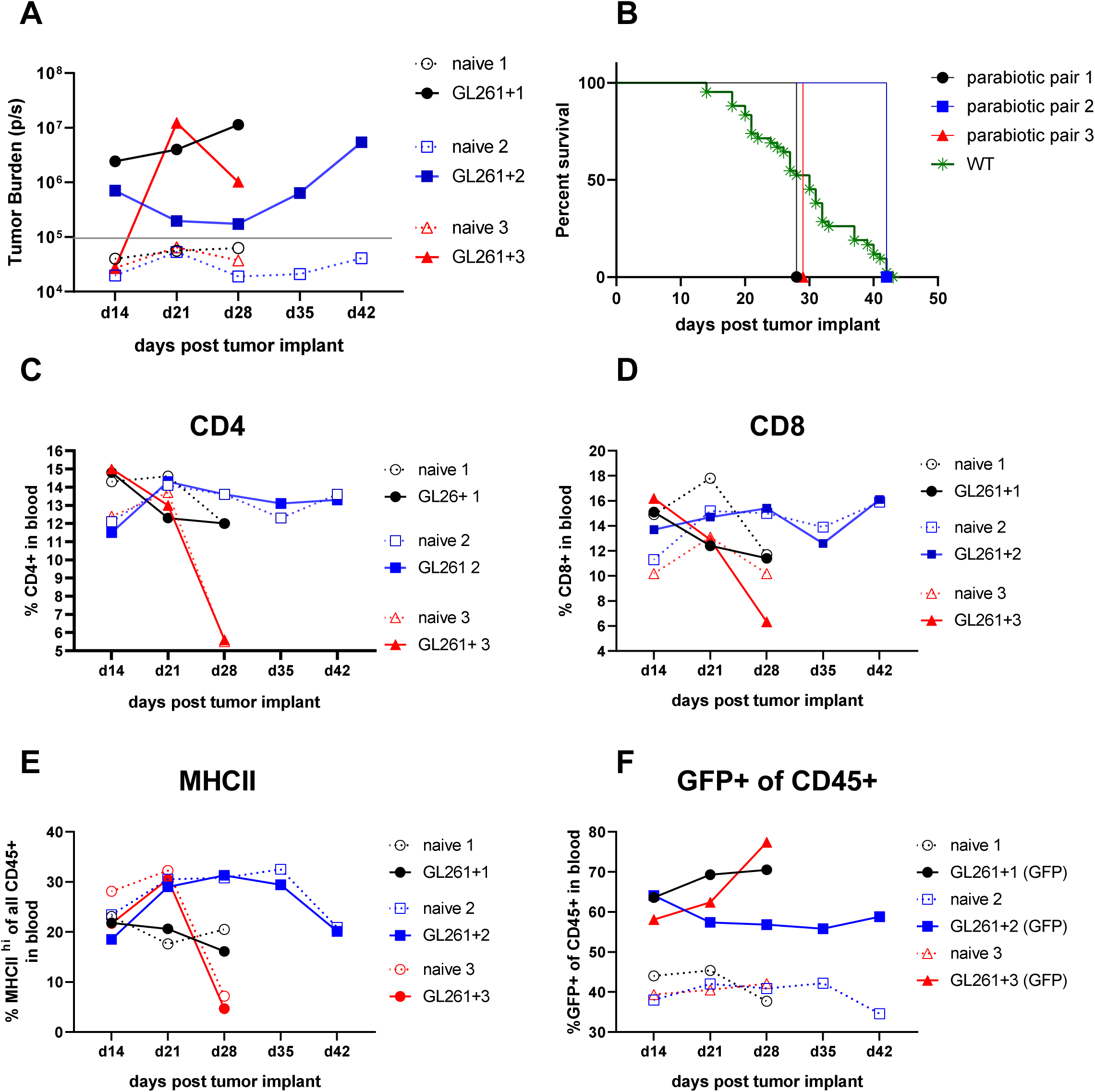
Immunosuppression during experimental GBM is transferrable to naïve mice via blood circulation: Tumor burden is shown in tumor-bearing parabiont and tumor negativity is confirmed in naïve partner (A). Values below 10^5^ are considered tumor negative. Survival of 3 parabiotic pairs was within the range of WT mice (n=43) (B). Frequencies of CD4 (C), CD8 (D), and expression levels of MHCII (E) are shown in all mice following analysis of the blood by flow cytometry. Frequencies of GFP^+^ cells were maintained within all 6 mice throughout the experiment (F).

**Figure S4:**
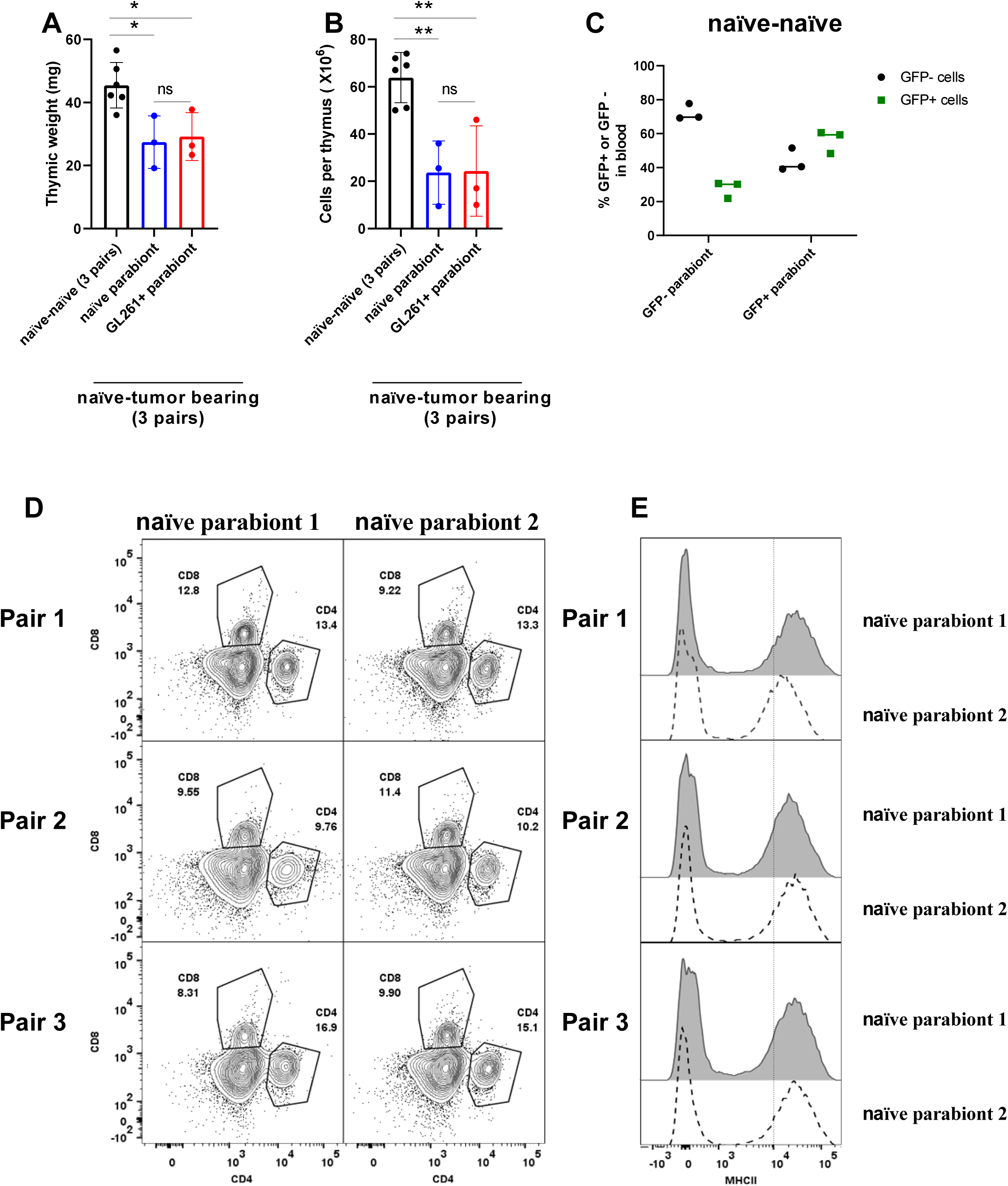
Hallmark features of immunosuppression do not occur following parabiosis of naïve to naïve mice: we performed parabiosis on 6 pairs of naïve mice. In each parabiotic pair, one mouse was GFP^+^. 30 days later, we implanted GL261-Luc cells in the brains of 3 mice. Naïve-tumor bearing parabionts were euthanized when tumor bearing mice became moribund. Naïve-naïve parabiont controls were euthanized on day 60 post parabiosis. Thymic weight (A) and cellularity (B) is compared between naïve-naïve and naïve-tumor bearing parabionts. Naïve to naïve mice had stable sharing of GFP+ cells (C). CD4, CD8 T cells (D), and MHCII expression on all CD45+ cells (E) in the blood do not decline in naïve-naïve pairs on day 60 post parabiosis.

**Figure S5:**
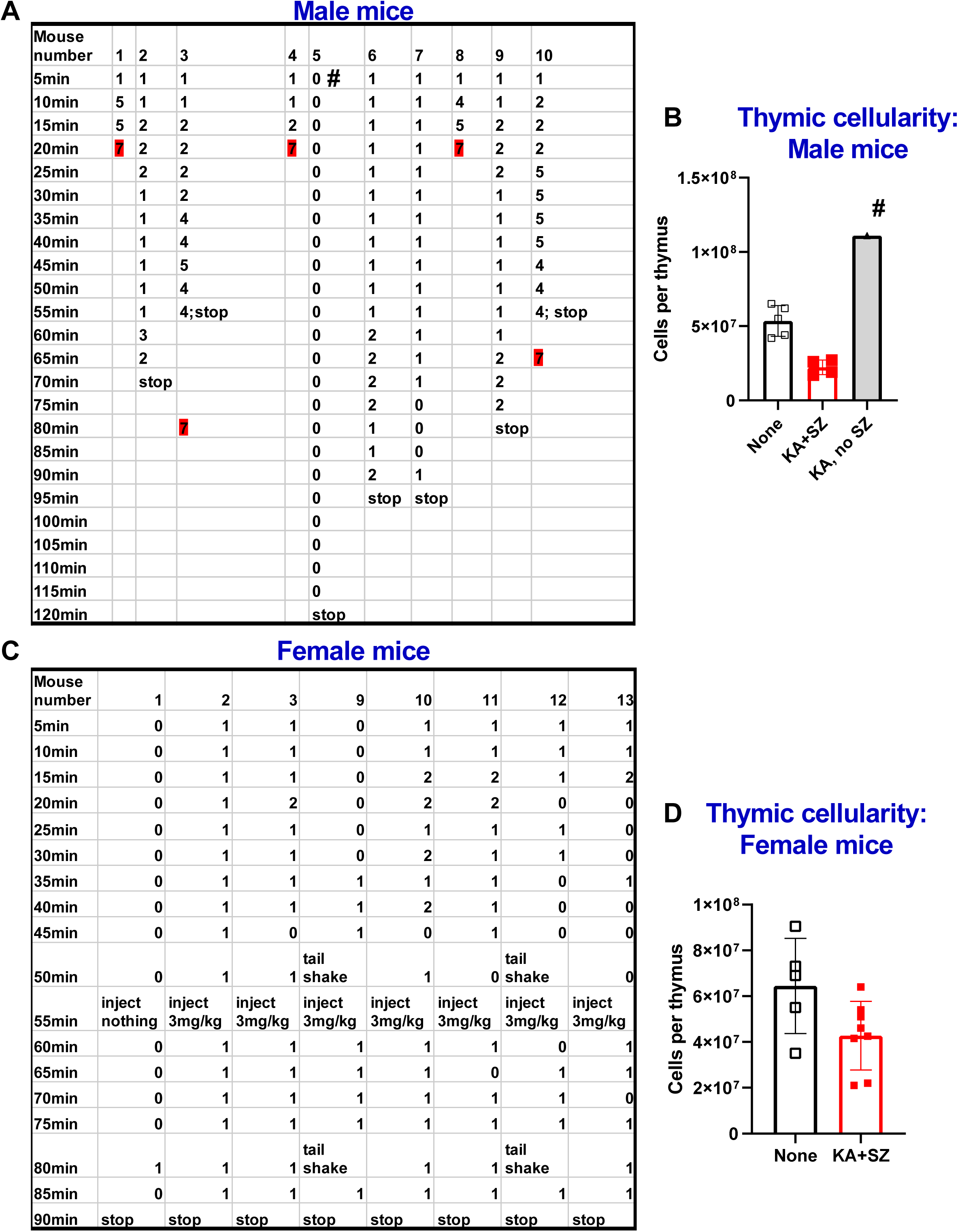
Seizure activity correlates with thymic involution on day 7 post KA injection: Individual seizure scores are shown for male (A) and female (C) mice as a function of time post KA injection. Thymic cellularity for male mice is quantified in (B). # denotes one mouse that did not have a seizure despite KA injection. This mouse did not develop thymic involution on day 7 post KA injections (B). Thymic cellularity in female mice is quantified in (D). These experiments were done in male mice and repeated in female mice in an independent experiment. N=4-8.

**Figure S6:**
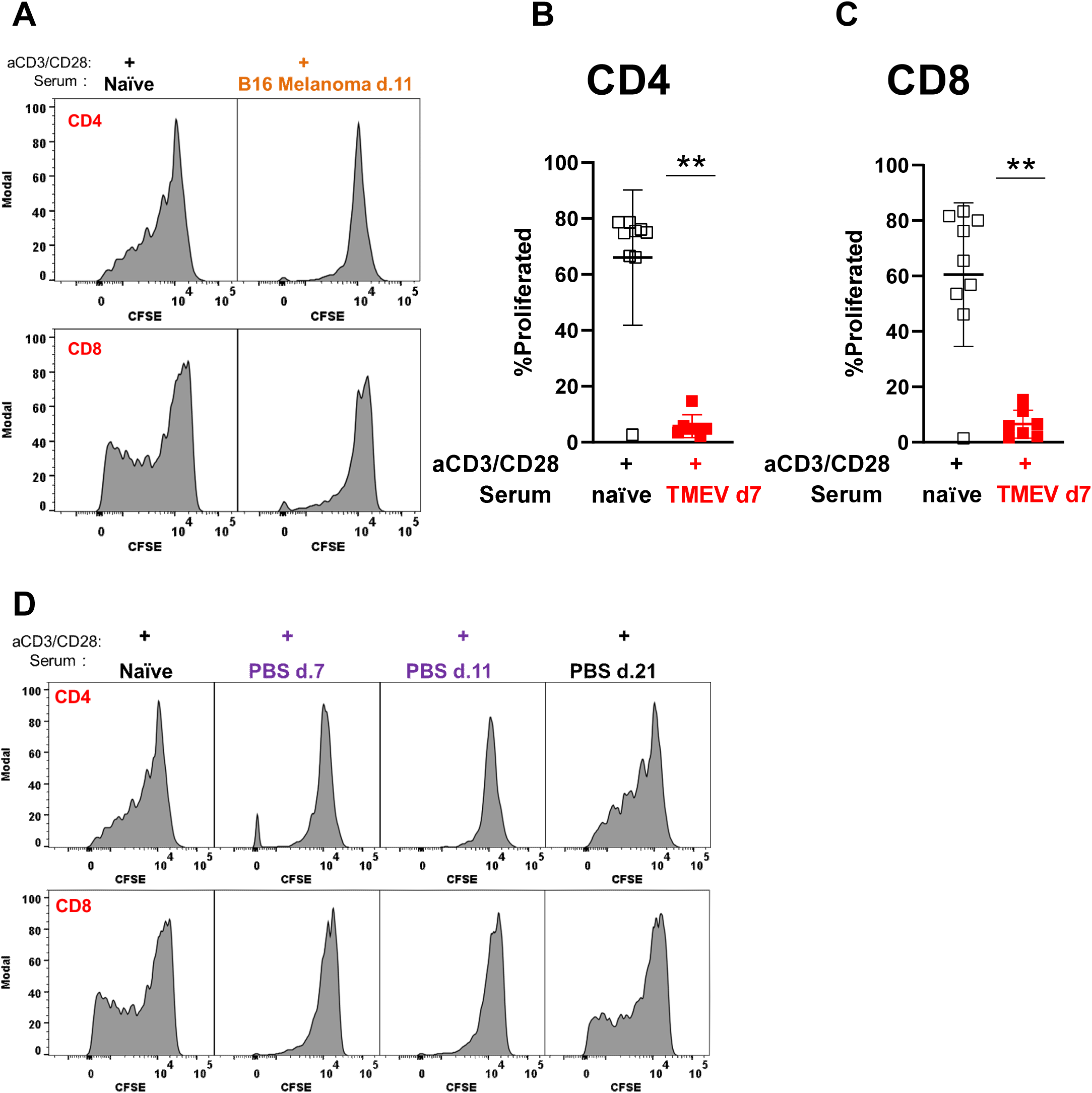
Serum obtained from mice with an ongoing neurological insult suppresses T cell proliferation *in vitro*: Representative histograms depict CD4 (top) and CD8 (bottom) T cell proliferation with anti-CD3/CD28 Dynabeads. Serum taken from mice injected with B16 melanoma (i.c) is tested for suppressive activity against T cells (A).Quantification of T cell proliferation following exposure to serum obtained from naïve or TMEV infected mice (day 7 post intracranial infection) is shown for CD4 and CD8 T cells (B-C). Serum obtained from PBS (i.c) injected mice on days 7, 11 and 21 post injection is tested for suppressive activity against T cells (D). These experiments were repeated twice with similar results. Representative histograms are shown.

**Figure S7:**
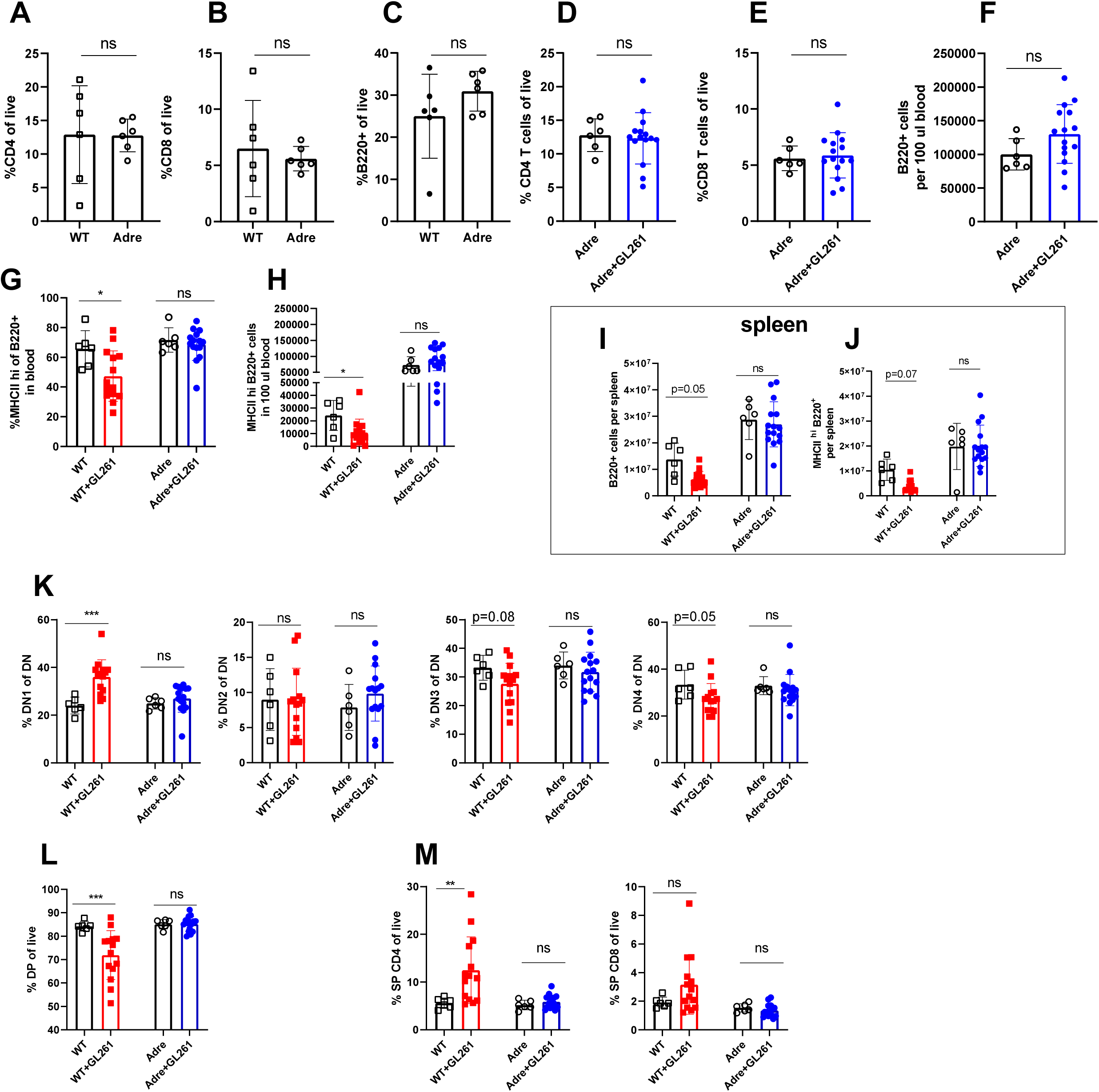
Hormones produced by the adrenal gland control immune organ size and cellularity at baseline: Frequencies of CD4 (A), CD8 (B), B220 (C) cells are compared in blood of naïve WT and adrenalectomized mice. Frequencies of CD4 and CD8 T cells in the blood do not change between naïve and glioma-bearing adrenalectomized mice (D-E). Numbers of B cells in 100 µl of blood does not change in glioma-bearing mice compared to naïve adrenalectomized controls (F). Frequencies and numbers of MHCII hi B cells declines in blood of glioma-bearing WT mice compared to WT controls, but do not change amongst adrenalectomized counterparts (G-H). Numbers of B cells (I), and numbers of MHCII hi B cells (J) are compared in spleens of WT and Adrenalectomized mice. Frequencies of DN1-DN4 (K), DP (L), and single positive CD4 and CD8 T cells (M) are shown for thymi isolated from naïve and glioma-bearing WT and adrenalectomized mice.

